# White matter reflects the childhood exposome

**DOI:** 10.64898/2026.05.04.722687

**Authors:** Briana Macedo, Joëlle Bagautdinova, Steven L. Meisler, Matthew Cieslak, Lucinda M. Sisk, Christos Davatazikos, Alexandre R. Franco, Meike D. Hettwer, Arielle S. Keller, Gregory Kiar, Audrey C. Luo, Allyson P. Mackey, Michael P. Milham, Tyler M. Moore, Valerie J. Sydnor, Kevin Y. Sun, Fang-Cheng Yeh, Ran Barzilay, Damien A. Fair, Russell T. Shinohara, Aaron Alexander-Bloch, Theodore D. Satterthwaite

**Author notes:** Address correspondence to: Theodore D. Satterthwaite.

## Abstract

The childhood environment is critical for brain development. However, most neuroimaging studies examine individual environmental measures (e.g., socioeconomic status) or a limited set of exposures, obscuring how the combination of complex, real-world exposures jointly influence brain development. Here we investigated how white matter shape and tissue properties are linked to the childhood exposome, a multidimensional measure capturing over 300 environmental exposures. Using multi-shell diffusion MRI from 8,183 children (ages 9-10) in the ABCD study, we quantified microstructural and macrostructural properties across 62 person-specific white matter tracts. The exposome showed widespread and highly replicable associations with both white matter microstructure and macrostructure: more advantaged environments were associated with larger tract macrostructure and lower orientation dispersion. Principal component analysis revealed that the dominant axis of exposome-white matter covariation aligns with the cortical sensorimotor-association hierarchy, such that tracts spanning this hierarchy exhibit the strongest associations with the exposome. Multivariate models demonstrated that patterns of white matter features explained 25% of the variance in the exposome in unseen individuals. Notably, white matter-based prediction of cognition was markedly reduced after accounting for the exposome (∼82% reduction in explained variance), indicating that brain-cognition associations overlap substantially with variance captured by the exposome. These findings generalized to independent data from the Healthy Brain Network (*n*=869), which differs substantially from ABCD in MRI acquisition, participant selection, and childhood environments. Together, these results suggest that white matter architecture strongly reflects the childhood environment.

## Introduction

The childhood environment plays a critical role in shaping brain development, with profound consequences for an individual’s lifelong cognition and mental health^1,2^. Emerging evidence suggests that environmental exposures in childhood may alter the pace of brain development^1–10^. Accelerated development is hypothesized to serve as an adaptation to early adversity^3,5,9,11^. However, this short-term adaptation may restrict sensitive periods of plasticity, with downstream consequences for brain function, psychopathology, and cognitive function^1,2,4,5^. In contrast, more enriched environments, often associated with higher socioeconomic status (SES), have been linked to more protracted development, potentially extending periods of plasticity and thus supporting gradual refinement of neural systems^4,6^. Accumulating evidence suggests that developmental plasticity ascends the cortical hierarchy defined by the sensorimotor-association (S-A) axis. The S-A axis ranges from unimodal, lower-order sensorimotor cortical regions to higher-order, multimodal association regions and reflects a core organizational gradient of cortical function and development^12^. Large-scale white matter (WM) tracts support long-range communication across this cortical hierarchy^13–16^ and the myelination of WM is critical for efficient signal conduction and network synchronization^16–18^. However, despite growing evidence that the environment shapes brain development, its influence on white matter structure remains incompletely understood. Specifically, it is unclear whether environmental effects are broadly distributed across white matter or instead preferentially impact long-range connections that traverse the cortical hierarchy. Because white matter tracts link regions that mature at different times, their developmental windows may span the combined periods of plasticity across connected regions, potentially extending their sensitivity to environmental influences. Here, we capitalize on both large-scale data resources and advanced analytic methods to investigate how the late childhood exposome shapes white matter tracts that link brain regions along the S-A cortical hierarchy.

Prior work has shown that multiple aspects of the childhood environment are associated with brain structure and organization across development—including SES^19–25^, pollution^26–33^, educational opportunity^34^, prenatal exposures^28,29,35,36^, adversity^3,37–43^, and even extreme temperatures^44^. However, a key limitation in this literature is that many prior studies have examined these aspects of the environment in isolation; this approach may fail to capture the additive effects of exposures^45–48^. The “exposome” summarizes an individual’s environmental exposures^45,49,50^. Exposome-based approaches have shown that composite environmental measures explain more variance in cognitive and mental health outcomes than commonly used indicators of SES such as household income or parental education^46,51,52^.

Diffusion MRI (dMRI) is the dominant method of non-invasively imaging WM in humans. dMRI allows characterization of WM microstructure, which describes tissue properties such as tract organization, axon packing, and myelination^18^. However, many of the most widely used dMRI measures (e.g., fractional anisotropy [FA], mean diffusivity [MD]) are biologically nonspecific and can reflect multiple processes, including myelination, axonal density, fiber orientation, and local cellularity^18^. Neurite Orientation Dispersion and Density Imaging (NODDI) addresses some of these limitations by applying a biophysical compartment model to multi-shell dMRI data, enabling estimation of microstructural features that capture distinct compartments with unique diffusion behaviors (e.g., hindered, restricted, and isotropic diffusion)^53–57^. NODDI models of microstructure have been shown to be more sensitive to neurodevelopment compared to measures from tensor models; they are also less sensitive to the confounding effects of head motion and image quality^53,58^. Furthermore, powerful methods now allow the delineation of person-specific WM tracts, facilitating the measurement of personalized tract macrostructure and geometry^59^. Whereas microstructural measures index local tissue properties, these macrostructural measures characterize larger scale tract-level anatomy: for example, their length, diameter, surface area, volume, and end region geometry. Prior work suggests that macrostructural features have distinct developmental trajectories compared to microstructural features^14^. Macrostructure measures of WM anatomy have been only sparsely explored, potentially providing novel and complementary information beyond microstructure alone^14,59,60^.

While prior exposome-neuroimaging studies have linked environmental exposures to cortical and subcortical structure and functional connectivity, far less is known about how the exposome relates to WM shape and microstructure in youth^21,46,52,61,62^. Although some existing literature has linked individual environmental exposures to WM, these prior studies have notable limits: small samples, a focus on simple tensor-based measures of WM microstructure, and very limited information on the role of WM macrostructure^62^. In the context of such limitations, prior work has yielded heterogenous results^23,38,54,62^, and it remains unclear whether environmental effects are widespread, organized according to known principles of brain organization, or extend to macrostructural features. Given the central role WM plays in efficient communication across the cortex, there is a critical gap in our understanding of how a broad set of environmental exposures across childhood relates to diverse WM properties across the brain.

Here we investigated how the childhood exposome is linked to WM microstructure and macrostructure at the transition from late childhood to early adolescence. We hypothesized that the exposome would show widespread associations across both micro-and macrostructural features, and that these associations would be systematically patterned along the S-A axis. Specifically, we predicted that there would be stronger effects in tracts that span the cortical hierarchy. Further, we predicted that associations between WM and the exposome would share substantial variance with any observed associations between WM properties and cognition. To test these hypotheses, we used dMRI data from 8,183 children in the Adolescent Brain Cognitive Development (ABCD) study and state-of-the-art tractometry approaches to derive person-specific WM micro-and macrostructural features. Finally, we replicated our results in the Healthy Brain Network (HBN, *n*=869) dataset^63^ to show that associations generalize to an independent cohort.

## Results

We investigated whether the childhood exposome is reflected in WM micro-and macrostructural properties in youth. To accomplish this, we used baseline diffusion MRI (dMRI) data from 8,183 children in the ABCD study^64^, split into demographically matched discovery (*n*=4,082) and replication samples (*n*=4,101; **Table 1**)^65^. We processed dMRI data (including microstructure and macrostructure measures, see **Supplementary Table 1**) using the state-of-the-art, open source dMRI pipelines QSIPrep^66^ and QSIRecon^67^ (**Figure 1**; see **Methods**). All dMRI features were harmonized using a cross-validated implementation of CovBat^68^ that protected covariates of interest but also prevented information leakage across the discovery and replication samples (see **Methods**). First, we characterized tract-wise associations between the exposome and WM properties using mass-univariate analyses. The exposome score primarily comprised socioeconomic factors spanning both household-level (e.g., income, parental education) and neighborhood-level indicators (e.g., geocoded poverty), along with measures of enrichment (e.g., extracurricular activities) and family environment (e.g., parental marital status; see **Supplementary Table 2** for exposome factor loadings). Higher exposome scores reflect more advantageous environments. We then evaluated the univariate results within the context of the sensorimotor-association axis. We next used multivariate machine learning models to evaluate the strength of associations between high-dimensional patterns of WM features and the exposome in unseen individuals. We then examined how links between the exposome and WM relate to individual differences in cognitive performance. Finally, we tested whether these results generalized to an independent cohort, the Healthy Brain Network (HBN, *n*=869).

**Figure 1.**
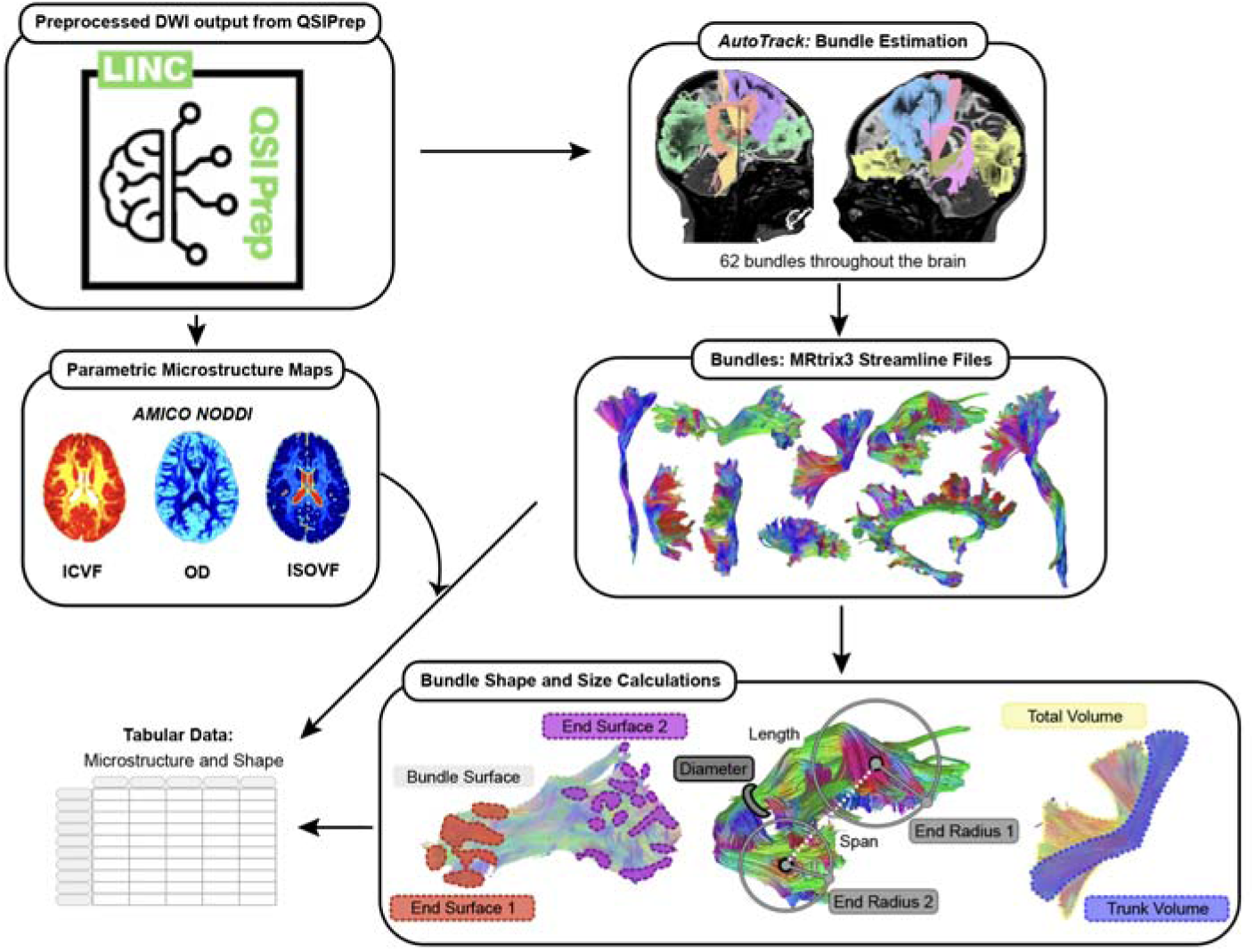
Quantification of person-specific tract macrostructure and microstructure. WM tracts were segmented at the individual level using AutoTrack. This yielded 62 tracts spanning the brain, which were exported as streamline files. For each tract, macrostructural properties were computed from tract geometry, including length, span, diameter, end radii, surface geometry, and trunk and total volume. Parametric microstructural maps derived from AMICO-NODDI (intracellular volume fraction [ICVF], orientation dispersion [OD], and isotropic volume fraction [ISOVF]) were generated. The median ICVF, OD, and ISOVF was calculated within each tract. Tract-wise macrostructural and microstructural measures were combined into a tabular dataset for analyses.

**Table 1.** Demographics of the ABCD and HBN samples. Discovery and replication samples from ABCD were generated using the ABCD Reproducible Matched Sample (ARMS), which were designed to balance participants across demographic features (including: site location, age, sex, ethnicity, grade, highest level of parental education, handedness, combined family income, and prior exposure to anesthesia)^65,69^. HBN is an independent, clinically enriched cohort used here for external validation.

| Sample | n | Age, mean (SD) | Female, n (%) | Male, n (%) |
| --- | --- | --- | --- | --- |
| ABCD Discovery | 4,082 | 9.98 (0.63) | 1,999 (49.0%) | 2,083 (51.0%) |
| ABCD Replication | 4,101 | 9.96 (0.63) | 1,922 (46.9%) | 2,179 (53.1%) |
| ABCD (Full) | 8,183 | 9.97 (0.63) | 3,921 (48.0%) | 4,262 (52.0%) |
| HBN | 869 | 9.81 (3.07) | 317 (36.5%) | 552 (63.5%) |

### Widespread, highly replicable associations between the exposome and white matter

We first asked whether the childhood exposome is broadly associated with WM micro-and macrostructural properties across major WM tracts. Using mass-univariate linear regressions (controlling for age, sex, and image quality; see **Methods**), we found that the general exposome was significantly associated with the majority of WM tract features after FDR correction (*q*<0.05) in both matched split halves of the data **(Figure 2a)**. Higher exposome scores—reflecting more advantageous environments (e.g., higher familial and neighborhood SES)—were linked to lower intracellular volume fraction (ICVF) and orientation dispersion (OD) across most tracts (**Figure 2a-b**). Associations with isotropic volume fraction (ISOVF) were more heterogeneous, with negative effects in association tracts and positive effects in projection tracts (**Figure 2a-b**). In contrast to the microstructural findings, the exposome showed strong positive associations with almost all macrostructural features, particularly tract length and span (**Figure 2a-b**). These tract-wise effect estimates were highly replicable across matched discovery and replication halves of the data (*r*=0.93; **Figure 2c**).

**Figure 2.**
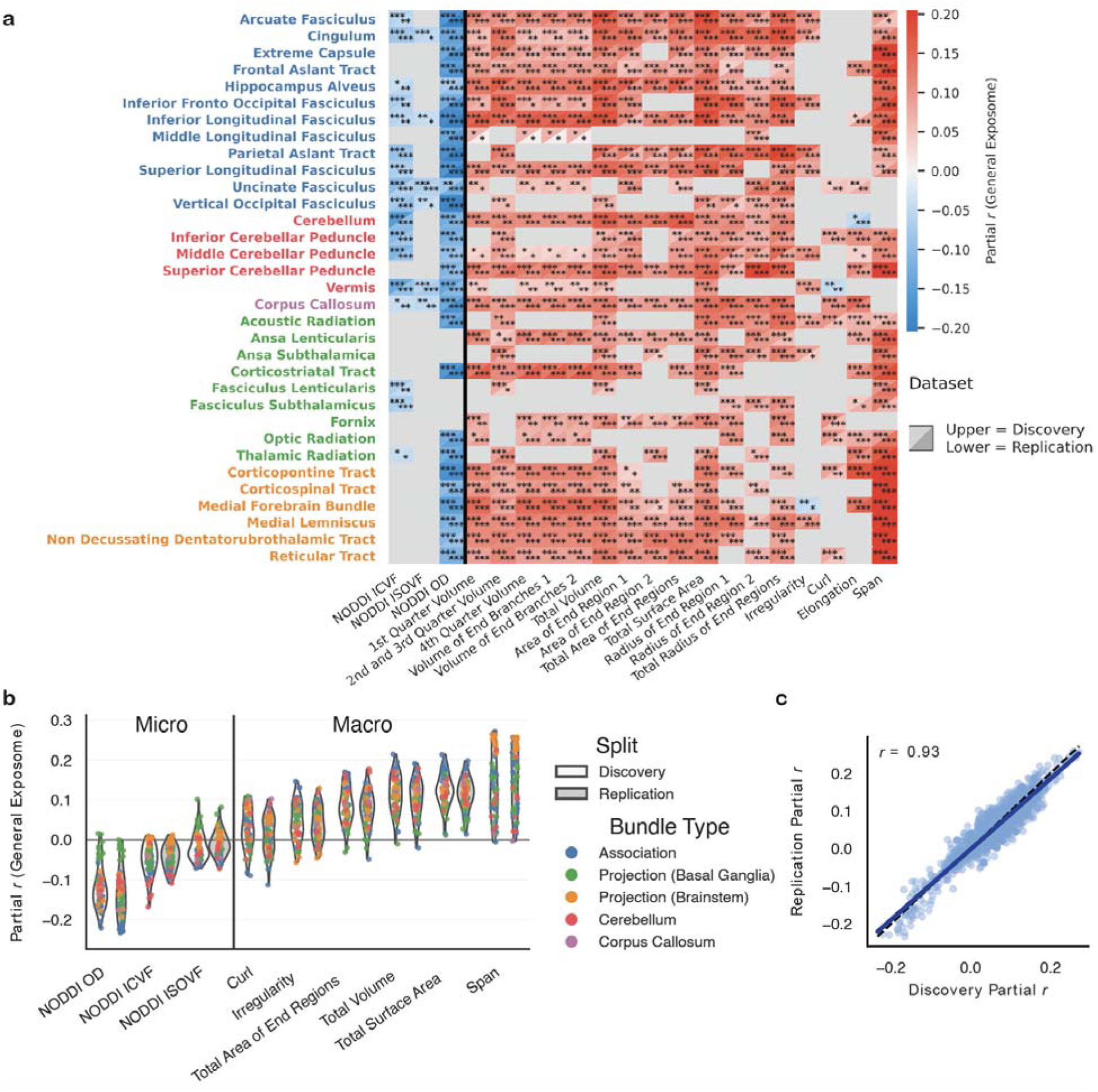
Widespread, highly replicable associations between white matter features and the exposome. **a)** Mass-univariate linear regressions testing the association between the exposome and each measure of tract microstructure and macrostructure. Metrics include three microstructure metrics defined by NODDI models and 18 measures of person-specific tract macrostructure. Each cell displays the signed square root of the exposome’s incremental signed effect size, computed as the change in model fit between a regression including the exposome score and an otherwise identical model with the exposome score removed. Effect sizes were averaged across hemispheres to simplify visualization. All models control for age, sex, and image quality. Asterisks denote FDR-corrected significance thresholds: * *q*<0.05, ** *q*<0.01, *** *q*<0.001. The upper triangle displays results from the discovery dataset, and the lower triangle displays results from the replication dataset. **b)** Violin plots summarizing the distribution of tract-wise effect sizes shown in panel a) for selected microstructural and macrostructural metrics. Each violin represents the distribution of effect sizes across WM tracts, with individual points colored by tract type (association, commissural, projection, and cerebellar tracts). Results are shown separately for discovery and replication samples, with similar effect size distributions across splits. **c)** Concordance of tract-wise effect sizes between discovery and replication samples. Each point represents a single association, plotted as the effect size from the discovery sample versus the replication sample. Concordance was evaluated using a Pearson correlation.

To assess the stability of these findings across modeling choices, we conducted several sensitivity analyses. A nearly identical pattern of effects emerged when repeating the mass-univariate analyses using diffusion kurtosis imaging (DKI)-derived microstructure metrics in place of NODDI-derived microstructure metrics (**Supplementary Figure 1**). Including total brain volume (TBV) as an additional covariate yielded broadly similar results, although effect sizes for macrostructural measures were attenuated and some associations no longer reached significance (**Supplementary Figure 2**). Importantly, tract-wise effect estimates from these models that corrected for TBV remained highly replicable across matched discovery and replication halves of the data (*r*=0.86). Finally, we found no evidence for sex-by-exposome interaction effects (**Supplementary Figure 3**).

### Principal components of white matter-exposome associations align with the cortical hierarchy defined by the S-A axis

Given the widespread and highly replicable nature of tract-wise exposome associations, we next interrogated how these effects are spatially organized across the brain. To summarize mass-univariate exposome-WM associations, we applied principal component analysis (PCA) to the tract-by-metric matrix of partial correlations (**Figure 2a**) averaged across discovery and replication splits. The first principal component (PC1) captured a dominant axis of covariation in exposome-WM associations across tracts (**Figure 3a**), explaining 35% of the variance in association patterns across tracts and metrics. Positive PC1 loadings were strongest for macrostructural features related to tract size including total volume, surface area, and tract length (**Figure 3a**). In contrast, as expected by the opposite sign of the mass-univariate associations, certain NODDI-derived microstructural measures loaded negatively onto this component, with the largest contribution from OD. Curl and elongation also showed negative loadings, though their contributions were relatively small. Each tract received a PC1 score, where higher values reflect stronger expression of the exposome association pattern observed in the mass-univariate analyses.

**Figure 3.**
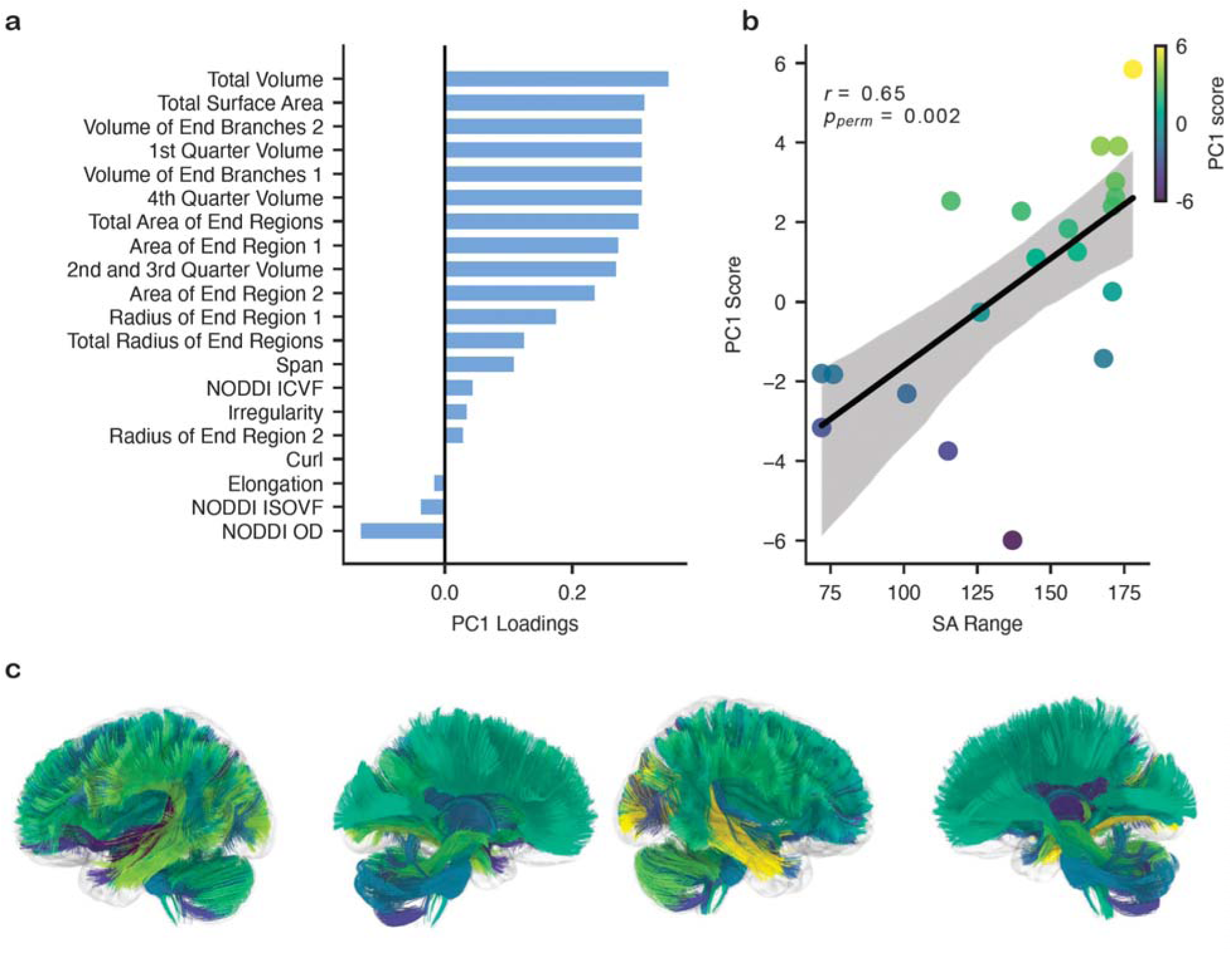
White matter-exposome associations vary systematically with span along the sensorimotor-association axis. We performed principal component analysis (PCA) of the tract-by-metric matrix of effect sizes obtained from mass-univariate linear regression models relating WM metrics to the general exposome score. Values were averaged across discovery and replication splits prior to PCA. **a)** Loadings of tract-level microstructure and macrostructure metrics from the first principal component (PC1) derived from this PCA. **b)** Association between tract PC1 scores and sensorimotor-association (SA) axis range (difference between the maximum and minimum SA rank of regions connected to each tract) across WM tracts. Each point represents a tract, colored by its PC1 score. The black line indicates the linear regression fit with 95% confidence interval. Pearson’s correlation coefficient (*r*) and corresponding *p*-value are shown. **c)** PC1 scores of tracts across the brain. Left hemisphere, lateral and medial slice; right hemisphere, lateral and medial slice.

We next asked whether this dominant axis of exposome-WM covariation was spatially organized along the hierarchical sensorimotor-association (S-A) axis. Given that WM tracts that span the S-A axis undergo protracted development, here we evaluated whether associations between the childhood exposome and WM properties varied by the S-A axis range (i.e., span) of WM tracts. We found that PC1 scores were positively associated with the S-A axis range of tracts (*r*=0.65, *p*=0.002; **Figure 3b**). Thus, tracts spanning a broader portion of the cortical hierarchy tended to have stronger associations with the exposome.

This relationship was preserved when using DKI-derived WM microstructure metrics instead of NODDI-derived metrics, suggesting that the result is robust to WM model choice (*r*=0.69, *p*=0.001; **Supplementary Figure 4**). We repeated this analysis separately in the discovery and replication splits, performing PCA within each split independently. Metric loadings were highly consistent across split halves (*r*=0.95), indicating a stable structure. PC1 scores were positively associated with the S-A axis in both splits (discovery: *r*=0.55, *p*=0.012; replication: *r*=0.67, *p*=0.001, **Supplementary Figure 5**). Together, these findings indicate that the dominant axis of exposome-WM covariation and its alignment with the S-A axis are replicable.

### Multivariate patterns of white matter features are strongly linked to the childhood exposome

Given the strong and robust univariate WM-exposome associations, we next evaluated multivariate patterns of association between WM features and the exposome in unseen data. Using demographically matched split halves of the ABCD sample (discovery and replication), multivariate ridge regression models were trained in one half of the data and tested in the other (and vice versa). Confound regression and harmonization was implemented to avoid information leakage across split halves. Performance was quantified as the correlation (*r*) between predicted and observed exposome scores in the held-out data. We trained separate models using microstructural features alone, macrostructural features alone, and their combination (the full feature set).

Models trained on NODDI microstructural features alone showed correlations of *r*=0.41 and *r*=0.40 in the discovery and replication samples, respectively (*p*<0.001 across split halves, **Fig. 4a**), whereas models trained on macrostructural features performed slightly better, with *r*=0.48 and *r*=0.49 in the discovery and replication splits, respectively (*p*<0.001 across split halves, **Fig. 4b**). However, it should be noted that macrostructural models included a larger number of features (18 metrics) than NODDI-based models (3 metrics). Despite this difference in model complexity, both feature sets independently showed excellent predictive performance in unseen data. Models incorporating both macrostructural and microstructural features performed best, with correlations between observed and predicted exposome scores of *r*=0.50 and *r*=0.51 across split halves (*p*<0.001 across split halves, **Figure 4c**). Predictive performance remained robust after adjustment for total brain volume, with only a modest attenuation of effect sizes (*r*=0.45, **Figure 4d**). Consistent with the mass-univariate analyses, multivariate feature importance closely aligned with tract-wise univariate effect sizes: Haufe-transformed weights strongly correlated with partial correlations across WM features (**Supplementary Figure 6**), indicating that the predictive signal is broadly distributed across the brain.

**Figure 4.**
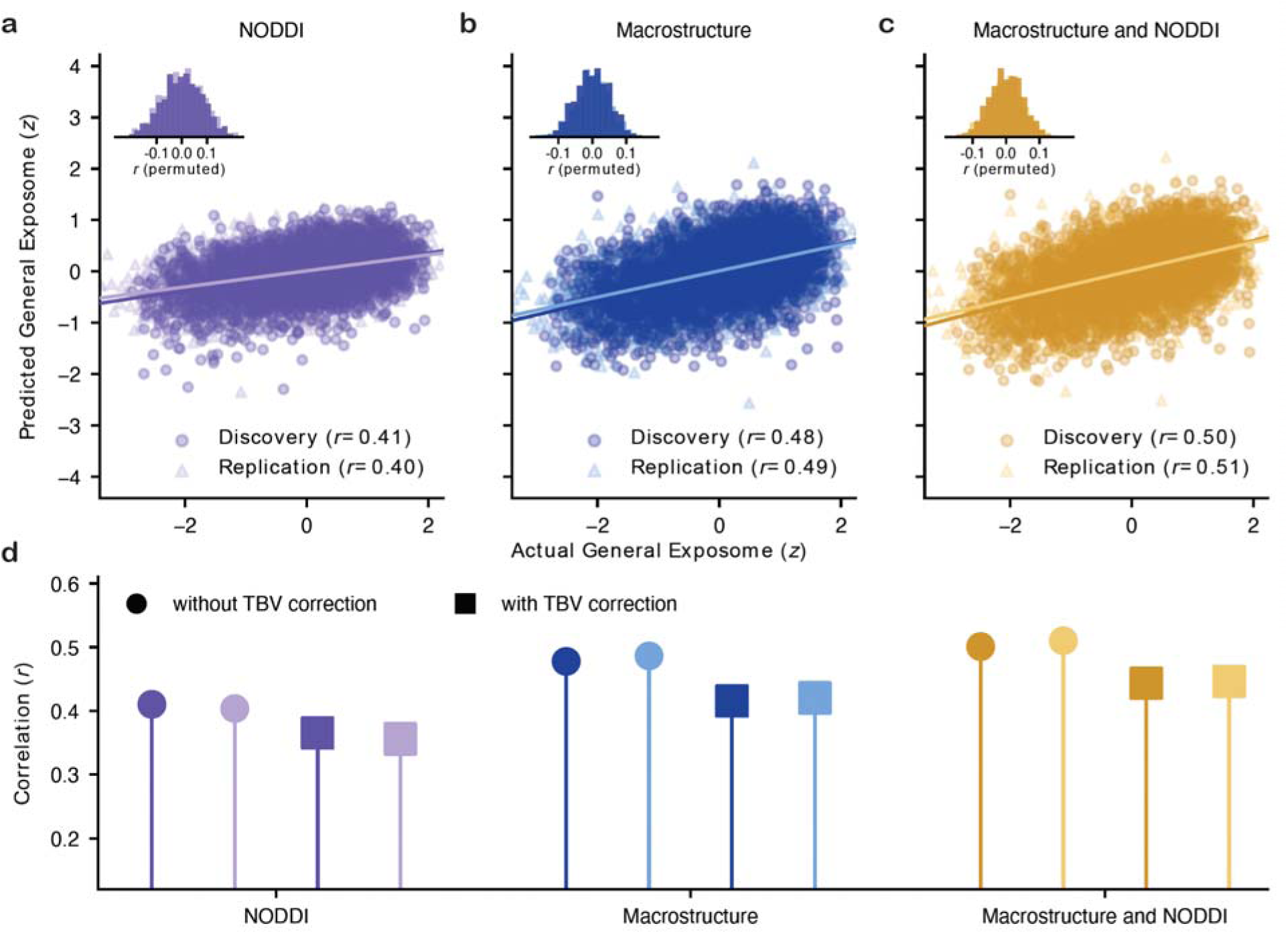
Robust multivariate associations between white matter features and the childhood exposome in unseen data. a-c) Association between observed and predicted exposome scores using two-fold cross-validated ridge regression across split halves. Each panel shows predicted versus observed exposome scores in unseen data for models trained using **a)** NODDI microstructural measures only, **b)** macrostructural measures only, and **c)** combined macrostructural and microstructural measures. Points represent individual participants from the discovery and replication split halves. Solid lines indicate the line of best fit estimated in held-out data. Insets show null distributions of prediction performance obtained through permutation testing. **d)** Pearson correlation (*r*) between observed and predicted exposome scores for each model, shown separately for analyses conducted with and without adjustment for total brain volume (TBV).

We observed similar prediction accuracy across a range of sensitivity analyses of the primary model incorporating both macrostructural and microstructural features. Comparable accuracy was observed when substituting nine DKI-derived microstructure metrics for the three NODDI-derived microstructure metrics (in combination with macrostructural features; *r*=0.51-0.53; **Supplementary Figure 7**) and when using partial least squares regression in place of ridge regression on the full set of features (*r*=0.49-0.50; **Supplementary Figure 8**).

### The exposome is more strongly linked to white matter than other commonly used measures

Given strong multivariate associations with the exposome (*r*=0.50-0.51; **Figure 4c**), we next evaluated whether the exposome provides added value over commonly used individual measures of the childhood environment. Using identical modeling and cross-validation procedures, WM features predicted the exposome substantially better than household income (*r*=0.36), parental education (*r*=0.28-0.30), and area deprivation index (ADI; *r*=0.37; **Supplementary Figure 9**). Despite lower associations for these individual environmental measures, feature weights were highly similar across models (**Supplementary Figure 10**), with strong correlations between Haufe-transformed weights derived from the exposome model and those derived from parental education (*r*_discovery_=0.88, *r*_replication_=0.93), income (*r*_discovery_=0.96, *r*_replication_=0.95), and ADI (*r*_discovery_=-0.90, *r*_replication_=-0.91) across folds. Together, these results highlight the value of using the exposome to summarize multiple aspects of the childhood environment.

### WM-cognition associations show substantial overlap with the exposome

Because the exposome is associated with general cognitive performance^52^, and environmental factors are known to influence both white matter and cognition^45,46,52^, we next examined whether WM-cognition associations in part reflect variance shared with the childhood environment. We first assessed associations between tract-wise metrics and general cognitive ability using a mass-univariate linear regression approach. Associations were modest but widespread, spanning both positive and negative effects (effect size range:-0.18 to 0.25; **Figure 5a**). Adjusting for the exposome reduced the magnitude of these tract-wise associations (adjusted *r* range:-0.07 to 0.14; **Figure 5a**). Feature-wise comparisons indicated that this attenuation reflects a consistent shrinkage of effect sizes within features, rather than tract-specific changes (**Figure 5b**). Adjustment reduced association strength across features in both the discovery and replication datasets, with an average decrease of ∼0.035 in |r| (≈43% reduction).

**Figure 5.**
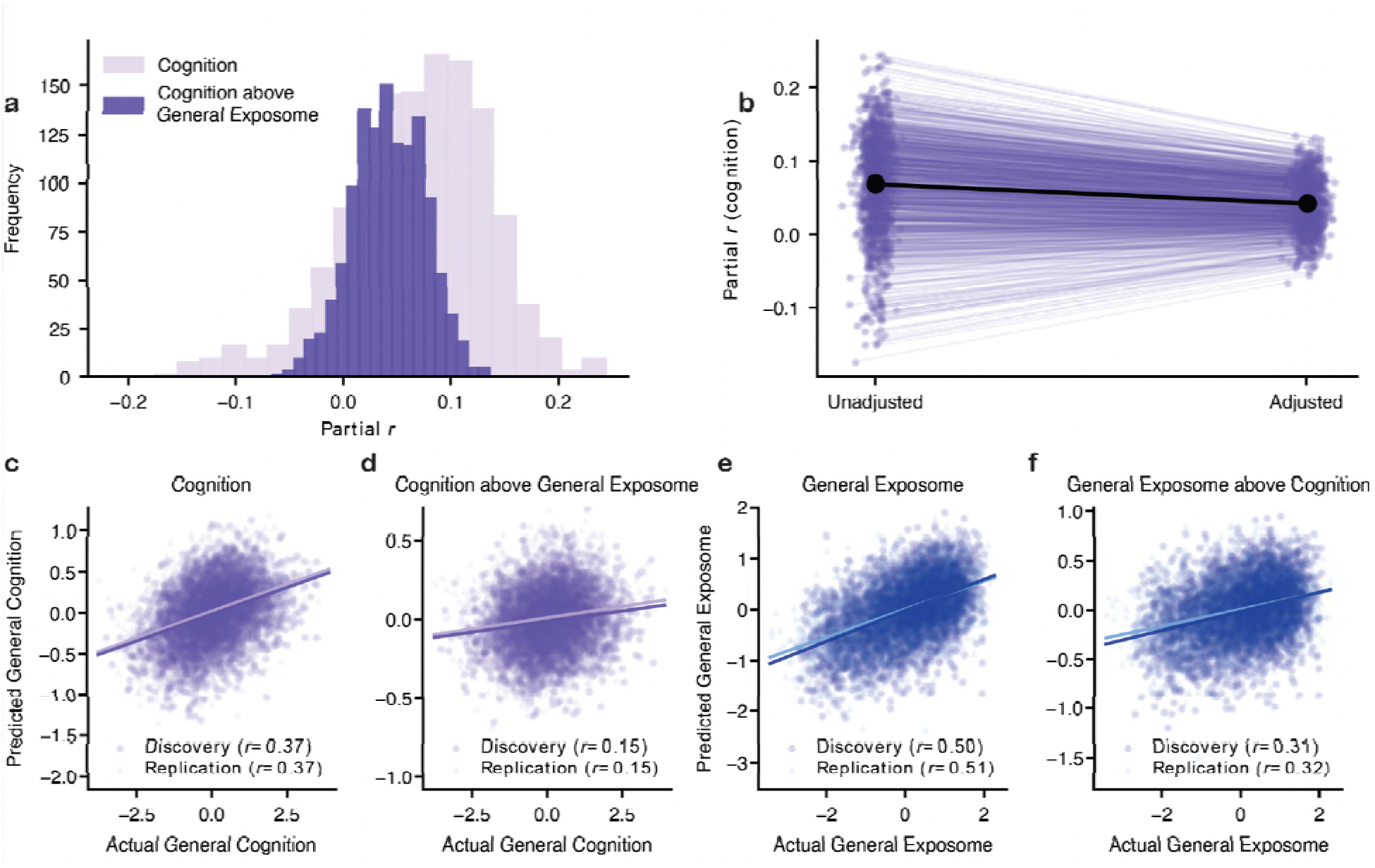
Associations between cognition and white matter share substantial variance with the exposome. **a)** Distribution of mass-univariate effect sizes between each tract-wise feature and general cognition, shown for models unadjusted (light) and adjusted (dark) for the general exposome score. **b)** Feature-wise correspondence between unadjusted and adjusted effect size values, showing decline in associations between WM and cognition when adjusting for the general exposome factor. **c)** Multivariate ridge regression model predicting general cognition: out-of-sample predicted vs. observed cognition scores (discovery and replication). **d)** Multivariate ridge regression predicting cognition after regressing out the exposome from cognition. **e)** Multivariate ridge regression predicting the exposome: out-of-sample predicted vs. observed exposome scores (discovery and replication). **f)** Multivariate ridge regression predicting the exposome after regressing out cognition.

We next extended these findings to the multivariate setting. WM features showed moderate out-of-sample associations with general cognition (*r*=0.37 across split halves, **Figure 5c**), corresponding to approximately 14% of variance explained. However, when the exposome was regressed from WM features prior to model fitting, association strength for cognition dropped substantially (*r*=0.15 across split halves; **Figure 5d**), explaining only 2-3% of variance, a relative reduction of 82%. This indicates that a substantial portion of WM-cognition covariance overlaps with variance captured by the exposome. In contrast, associations between WM features and the exposome were stronger and more robust to adjustment. WM features showed stronger out-of-sample associations with the exposome than with cognition (*r*=0.50 in discovery, *r*=0.51 in replication; **Figure 5e**). After regressing out cognition, associations remained fairly robust (*r*=0.31 in discovery, *r*=0.32 in replication; **Figure 5f**).

When total brain volume was added as a covariate, a similar pattern emerged, although effect sizes were even more attenuated: WM-cognition associations decreased to *r*=0.27-0.28, and additional adjustment for the exposome reduced associations to *r*=0.10 **(Supplementary Figure 11)**. In contrast, WM-exposome associations decreased to *r*=0.44, but remained relatively robust after adjustment for cognition (*r*=0.29-0.30).

### White matter-exposome associations generalize to an independent cohort

To assess whether exposome-WM associations generalized beyond the ABCD cohort, we evaluated data from the Healthy Brain Network (HBN), a clinically enriched sample that differs substantially from ABCD in ascertainment strategy, age range, sex distribution, exposome variables, and imaging acquisition (**Table 1**). First, we examined whether mass-univariate associations between exposome scores and tract-wise features generalized across cohorts. Consistent with ABCD (**Figure 2a**), HBN showed a similar pattern of associations, with microstructural metrics (e.g., NODDI-derived measures) having negative associations with the exposome, and macrostructural features (e.g., volume and length) having positive associations (**Supplementary Figure 12**). While effect sizes were larger in ABCD, associations between individual WM features and the exposome were highly similar across datasets (averaged across split halves, *r*=0.83; **Figure 6a**).

**Figure 6.**
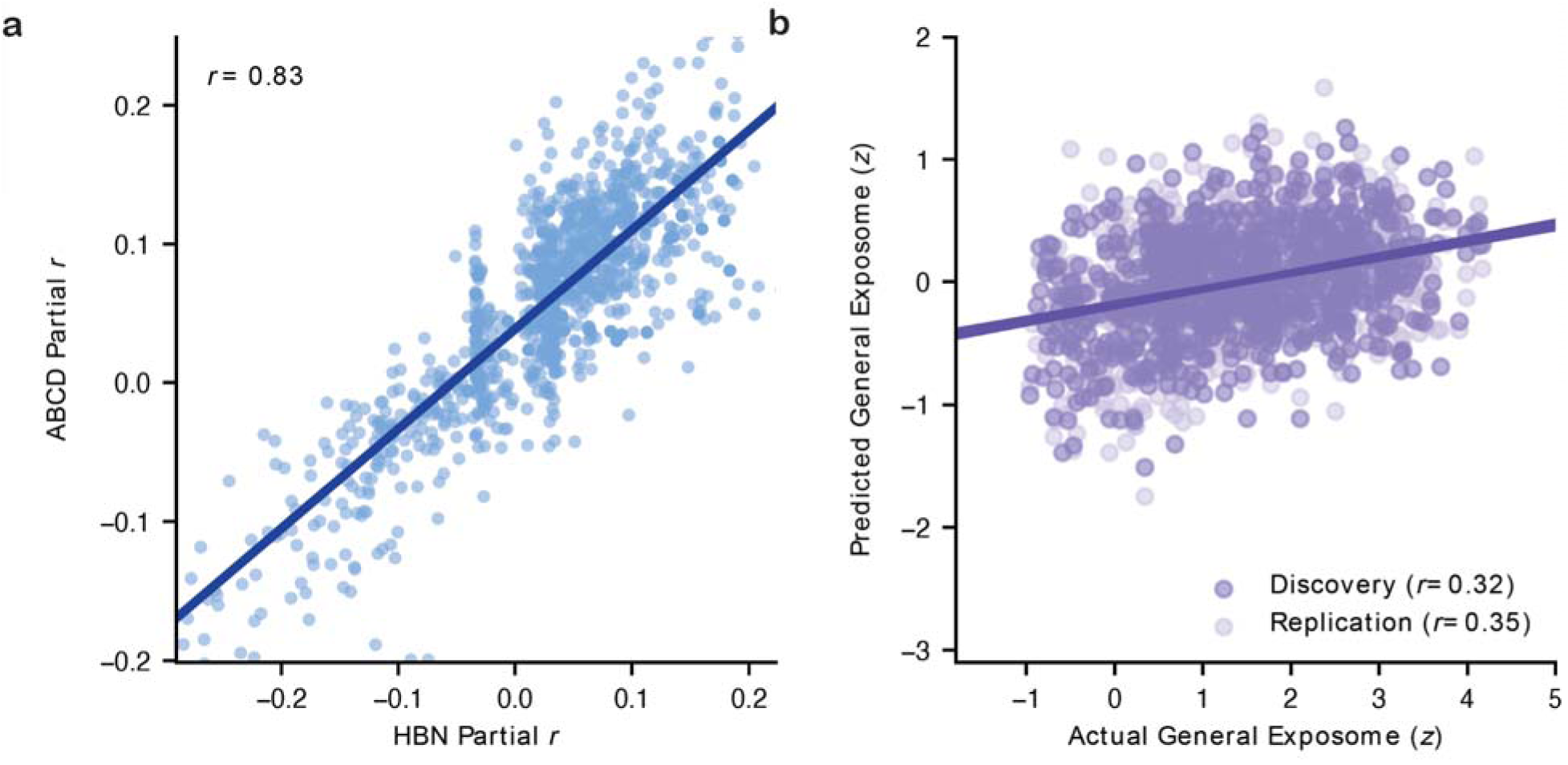
Results generalize to independent data from the Healthy Brain Network. **a)** Exposome-WM associations replicate across cohorts at the feature level. Each point represents effect size of the exposome with a tract-wise WM feature. The plot compares effect size values in HBN against the corresponding ABCD effect sizes, averaged across the two ABCD split halves. Strong concordance indicates robust replication of exposome-WM effects across independent datasets. **b)** Multivariate models trained to predict exposome scores from WM features in two matched ABCD split halves show consistent out-of-sample prediction when applied to unseen data from HBN without any additional tuning. The scatterplot shows observed versus predicted exposome scores in HBN for models trained in the ABCD discovery (dark purple) and replication (light purple) samples. Solid lines indicate best-fitting linear relationships and Pearson correlations (*r*) quantifying generalization performance (*p*<0.01 across split halves).

As a final step, we evaluated whether multivariate models trained on ABCD data could predict exposome scores in HBN participants. Ridge regression models trained separately in the ABCD discovery and replication samples showed significant prediction when applied to unseen participants in HBN without further tuning. Correlations between observed and predicted exposome scores were *r*=0.32 and *r*=0.35, using models trained on the ABCD discovery and replication cohorts, respectively (*p*<0.001 across split halves, **Figure 6b**). While prediction accuracy was lower than within-cohort performance in ABCD, these results emphasize that the pattern of associations between person-specific WM features and the exposome is broadly generalizable.

## Discussion

This study demonstrates that the exposome is associated with widespread and highly replicable differences in WM micro-and macrostructure in youth. Greater environmental advantage was associated with larger WM macrostructure and lower neurite orientation dispersion (OD). Effects were consistent across independent samples and analytic approaches. Exposome-WM associations were strongly and positively aligned with a known hierarchical organizing principle in the brain: the sensorimotor-association (S-A) axis^12^. Multivariate models further demonstrated that WM features explained over 25% of the variance in the exposome in unseen individuals. Furthermore, we found considerable shared variance attributable to environmental factors in the associations between WM and cognition. These findings generalized to an independent dataset capturing a distinct sample with different demographic and clinical characteristics. Together, these results show that variation in childhood environment is reflected in white matter by late childhood.

Exposome-WM associations were evident across both microstructural and macrostructural features, indicating that environmental exposures relate to multiple scales of white matter organization. These findings extend prior work connecting specific environmental exposures, such as socioeconomic disadvantage, air pollution, and childhood adversity^3,10,23,25,28,31,39,41–43,70^, to differences in WM development, which has largely relied on tensor-based metrics such as fractional anisotropy and mean diffusivity^3,10,10,25,28,31,39,41,70^. In contrast, our findings show that cumulative environmental influences are associated with widespread differences in white matter organization that are not fully captured by single indicators of socioeconomic context, including household income, parental education, or area deprivation index. By integrating sensitive microstructural metrics^53^ with person-specific macrostructural features and a multidimensional exposome measure, this work provides a more complete picture of how multiple environmental exposures relate to white matter properties. This approach revealed strong, consistent, and replicable associations between WM features and exposome scores, above and beyond effect sizes seen in smaller studies investigating individual associations (e.g., correlations |*r*| often <0.1).

We observed robust associations between the exposome and WM macrostructure. Although microstructure features have been extensively studied in the developmental literature, macrostructure measures^59^ that go beyond conventional volume metrics to characterize tract geometry (e.g., length, surface area, and end-region morphology) remain only sparsely explored. We found that WM tract macrostructure properties had strongly positive associations with the exposome and significantly predicted the exposome in unseen individuals. Lifespan studies show that many WM macrostructural features (e.g., volume, length, and surface area) increase through childhood into early adulthood, with particularly protracted development in tracts connecting to the prefrontal cortex^14^. These trajectories appear coordinated with microstructural development and vary by cortical target^14^. Prior work suggests that macrostructural features related to tract endpoint geometry show substantial between-subject variability^51^. Furthermore, combining macrostructural measures with microstructure can improve prediction of behavioral phenotypes such as cognition^71^. Our work extends this literature, showing that tract macrostructure is strongly associated with childhood environmental exposures captured by the exposome. One possible explanation is that macrostructural features such as tract volume, length, and geometry reflect white matter growth and refinement. Notably, this remained true even when accounting for total brain size, suggesting that they are not simply driven by global differences in brain size across participants. These findings raise the possibility that WM macrostructure represents a pathway through which environmental exposures shape large-scale brain communication during development.

We also observed robust associations between the exposome and WM microstructure. Developmental studies using NODDI measures of WM microstructure have shown that intracellular volume fraction (ICVF) increases in large-scale WM tracts throughout childhood and adolescence, reflecting increased neurite density due to axonal packing and myelination^14,53,54,58,72–75^. Orientation dispersion (OD), which reflects the angular variation of neurite fibers (e.g., parallel vs. fanning or crossing fibers), provides complementary information about white matter architecture. We found that more disadvantaged environments were associated with higher ICVF, particularly in association and cerebellar tracts, as well as higher OD across tracts, indicating more complex fiber organization. These patterns may reflect relatively more mature or altered microstructural organization in these pathways. One potential interpretation of these results is that environmental exposures may impact the pace of WM development. This is in line with a growing body of work suggesting that the childhood environment may influence the pace of neurodevelopment^1,4–8^, which may be adaptive in the short term, but may have consequences for cognition and mental health^1,2,4,5^. Notably, much of the literature on the influence of the environment on the pace of neurodevelopment has focused on gray matter in cortical^9,41^ and subcortical structures^5,9,76^, leaving a gap in our understanding of how the environment may impact WM development^4^. Given the strong effects we observe here in children ages 9-11 years, our findings are consistent with the possibility that environmental exposures influence the developmental timing of WM at early ages^77–79^. An alternative interpretation is that environmental disadvantage is associated with stable differences in WM microstructure across development, rather than differences in developmental timing. Distinguishing between these possibilities represents an important direction for future research and will require longitudinal data spanning earlier developmental periods.

We found that the childhood exposome was differentially related to WM tract measures based on the extent to which they span the cortical hierarchy, with stronger effects in tracts linking sensorimotor and association regions. Converging evidence indicates that brain development unfolds hierarchically across development, progressing from unimodal sensorimotor regions to higher-order association cortices along the S-A axis^12^. This developmental progression along the S-A axis holds across diverse structural and functional brain features, including cortical volume^80^, intrinsic activity^81^, functional connectivity^82,83^, cortical myelination^84^, structure-function coupling^85^, and thalamocortical structural connectivity^86^. WM development also aligns with the S-A axis: tract segments near association cortices show later and more protracted development compared to regions near sensorimotor cortices, paralleling the prolonged maturation of association cortex itself^87^. Furthermore, recent work demonstrated that tracts which span a broad range of the cortical hierarchy—i.e., connecting lower-order sensorimotor to higher-order association cortices—exhibit greater associations with age^15^. Our work extends this literature, revealing that tracts spanning the cortical hierarchy have the strongest associations with the exposome. These tracts link lower-order sensorimotor regions with higher-order association cortex, thus connecting cortical regions that support a diverse range of cognitive functions^15,88,89^. The heightened environmental sensitivity of WM tracts that bridge sensorimotor and association systems may reflect their more protracted development and their role in linking regions with distinct functions and developmental timelines and may contribute to environmentally driven differences in higher-order cognition.

WM supports cognition by connecting distributed cortical regions^89–91^. The childhood environment is strongly associated with neurocognitive development^37,88,92^. A key aspect of the childhood exposome, socioeconomic status, has been shown to shape both WM development and cognitive outcomes^46,52,93^. Extending this literature, we found that associations between WM and cognition were markedly attenuated after accounting for the exposome. These results suggest that a substantial portion of WM-cognition associations in late childhood reflect variance shared with the childhood environment. These findings align with results from a recent brain-wide association study showing that socioeconomic status had the strongest associations with brain measures and explained much of the brain-behavior variance often attributed to cognition^70^. Consistent with this, neighborhood and familial socioeconomic disadvantage have been linked to WM microstructural features in children, with WM integrity potentially mediating the link between SES and cognitive performance^25^. Specifically, growing up in enriching environments such as neighborhoods with more green space has been associated with increased WM integrity and greater fluid and crystallized intelligence^94^. However, our analyses are cross-sectional and cannot resolve causal structure. The attenuation we observe after adjusting for the exposome could reflect several possibilities. First, our results are consistent with WM features serving as a mediating variable between the observed relationship between SES and cognition. Alternatively, these results are consistent with the impact of shared upstream factors that influence both white matter and cognition. Distinguishing between these and other explanations would require longitudinal data in early life (before age 10). In addition, the general exposome factor is heavily weighted toward socioeconomic and enrichment-related variables. Part of the overlap we observe may reflect shared variance related to educational opportunity and language environment.

A notable strength of the present study is that exposome-WM associations generalized to the Healthy Brain Network (HBN), an independent cohort that differs substantially from ABCD in ascertainment strategy, age range, sex distribution, clinical composition, imaging acquisition, and exposome measurement^63^. Importantly, HBN is a clinically enriched sample recruited through community advertisement targeting families with concerns about psychiatric symptoms^63^, resulting in substantially higher levels of psychopathology compared to ABCD, which was designed to be representative of the general population^64^. Furthermore, whereas the ABCD exposome was derived from a broad mixture of individual-, caretaker-, and neighborhood-level exposures, the HBN exposome was based exclusively on geocoded neighborhood-level indicators^52,92^. Despite these differences, the overall pattern of associations was similar across cohorts: mass-univariate tract-wise effect sizes were highly concordant across HBN and ABCD, and multivariate models trained in ABCD successfully predicted exposome scores in HBN participants. Although predictive performance was lower in HBN than within ABCD, this is expected given the marked differences between cohorts, including both the clinically enriched sampling strategy, different dMRI acquisitions, and the more restricted, neighborhood-only exposome measure in HBN. Together, these findings indicate that exposome-WM associations generalize across populations that differ not only in demographic and clinical characteristics, but also in how environmental exposures are defined and measured. In a field where brain-wide association studies have faced challenges related to reproducibility and inflated effect sizes in smaller samples^69,95–97^, this cross-cohort generalization provides especially strong support for the robustness of our results.

Our results should be considered in the context of several limitations. First, we used cross-sectional baseline data from the ABCD study to establish foundational associations and lay the groundwork for future studies investigating the exposome and WM. However, this design limits inferences about within-individual development and restricts interpretation to the studied age (9-10). While we identify widespread and spatially organized associations between the childhood exposome and WM properties, these results cannot causally determine whether the exposome is related to differences in the timing or pace of WM maturation. Notably, these associations are already evident at ages 9-10 years, indicating that exposome-related differences in WM are established by late childhood. Because the ABCD study begins at ages 9-10, we are unable to capture earlier developmental periods during which exposome-related influences on WM may begin to emerge. Second, the exposome is a composite measure of many different environmental factors, limiting our ability to identify which specific components are most directly related to WM. Nevertheless, our sensitivity analyses revealed weaker associations between WM and single-measure household socioeconomic indicators, highlighting the relevance of the exposome approach in capturing the complex influence of the environment on WM organization. This is a core tradeoff of the exposome approach, which captures multidimensional effects across collinear exposures at the expense of insight into the precise effects of a single given variable. Finally, these results do not differentiate potential gene-by-environment effects; variables contributing to the exposome may have heritable aspects^98^. Future work leveraging twin data in ABCD could contribute to disentangling genetic and environmental contributions^99^.

The findings of this study point to several important avenues for future research. The large amount of variance in WM explained by the exposome may reflect the extent to which children’s environments in the United States are characterized by significant structural inequality^100–104^. Whether similar patterns would be observed in settings with a different range or disparate forms of inequality remains to be explored. Importantly, our findings suggest that environmental influences on WM are already detectable in late childhood, implying that these effects may become embedded even before age nine. These results set the stage for future longitudinal investigations earlier in life that can provide insight into when exactly environmental exposures become embedded in the developing WM, given evidence that environmental exposures can have especially strong effects during sensitive developmental windows^10,19,38^. Furthermore, our work highlights the importance of developing early interventions that support child development to target resiliency factors that protect from adversity during sensitive periods and to modify environmental conditions themselves, promoting better cognitive and mental health outcomes^62,92,105^. Finally, animal studies will be essential to determine the causal mechanisms underlying environmental influences on WM development and to clarify when during development these environmental effects become embedded in WM structure. Together, these findings suggest that the childhood environment is strongly associated with WM organization. More broadly, they raise the possibility that the childhood environment becomes biologically embedded in WM architecture, with potential downstream consequences for cognition and mental health^5,29,38,44^.

## Methods

### Dataset and Participants

We analyzed cross-sectional data from 8,183 children (ages 9-10) from the Adolescent Brain Cognitive Development (ABCD) Study^64^. Baseline diffusion MRI (dMRI) data were obtained from the ABCD-BIDS Community Collection (ABCC; release 3.1.0), available via the NIH Brain Development Cohorts (NBDC) Data Hub. All analyses were restricted to participants with complete diffusion imaging, environmental, and covariate data. Participants were assigned to discovery (*n*=4,082) and replication (*n*=4,101) samples using ABCD reproducible matched samples^65^, yielding two demographically matched subsets for split half analyses.

### Diffusion MRI Processing

Diffusion MRI preprocessing and reconstruction followed Meisler et al. (2026)^58^; we summarize relevant image processing here. All T1w, dMRI, and fieldmap images were preprocessed with the BIDS application^106^ *QSIPrep*^66^ version 0.21.4, which is based off of *Nipype* version 1.8.6^107,108^ and uses *Nilearn*^109^ version 0.10.1 and *Dipy*^110^ version 1.5.0. Data were postprocessed with *QSIRecon*^111^ version 1.0.0rc2, which is based off of *Nipype* 1.9.1 and uses *Nilearn* version 0.10.1 and *Dipy* version 1.8.0. The following paragraphs contain partially-edited text generated by *QSIPrep* and *QSIRecon*, which is distributed with a Creative Commons Zero (CC0) license with the express intention of being included in manuscripts.

#### Anatomical preprocessing

The T1w image was corrected for intensity non-uniformity using *N4BiasFieldCorrection*^112^ from *ANTs*^113,114^ version 2.4.3, and used as an anatomical reference throughout the workflow. The anatomical reference image was reoriented into AC-PC alignment via a rigid transformation extracted from a full affine registration to the MNI152NLin2009cAsym template. A full nonlinear registration to the template from the AC-PC space was estimated via symmetric nonlinear registration (SyN) using *antsRegistration*. Brain extraction was performed on the T1w image using *SynthStrip*^115^, and automated segmentation was performed using *SynthSeg*^116,117^ from *FreeSurfer*^118^ version 7.3.1.

### Diffusion MRI preprocessing

MP-PCA denoising as implemented in *MRtrix3*’s^119^ *dwidenoise*^120^ was applied with a 5-voxel window. After MP-PCA, Gibbs unringing was performed using *TORTOISE*’s^121^ *Gibbs*^122^. Following unringing, the mean intensity of the dMRI series was adjusted so all the mean intensity of the *b* = 0 images matched across each separate dMRI scanning sequence. *FSL*’s^123^ (version 6.0.8) *eddy* was used for head motion correction and eddy current correction^124^. *eddy* was configured with a *q*-space smoothing factor of 10, a total of 5 iterations, and 1000 voxels used to estimate hyperparameters. *eddy*’s outlier replacement was run^125^. *FSL*’s *TOPUP*^126^ was used to estimate a susceptibility-induced off-resonance field based on the reverse phase-encoded *b* = 0 reference image. The *TOPUP*-estimated fieldmap was incorporated into the eddy current and head motion correction interpolation. Final interpolation was performed using the Jacobian modulation method. The dMRI time series was rigidly aligned to AC-PC orientation to match the T1w image, preserving the original 1.7mm isotropic voxel resolution. B1 field inhomogeneity was corrected using *dwibiascorrect* from *MRtrix3* with the N4^112^ algorithm after corrected images were resampled.

#### Diffusion MRI postprocessing

dMRI reconstruction models were run as part of *QSIRecon*, resulting in a broad set of derived scalar maps. All scalar maps described below were produced in participant-specific DWI space (AC-PC orientation), as well as in MNI152NLin2009cAsym space (using nearest neighbor interpolation).

#### Neurite orientation dispersion and density imaging

Neurite Orientation Dispersion and Density Imaging (NODDI)^127^ is a biophysical model that characterizes microstructural features of brain tissue by estimating the relative contributions of intra-neurite, extra-neurite, and isotropic (CSF-like) compartments. The model provides indices such as intracellular volume fraction (ICVF, synonymous with neurite density index or NDI), orientation dispersion index (OD), and isotropic volume fraction (ISO), which reflect neurite density, angular dispersion, and free-water content, respectively. We used the Accelerated Microstructure Imaging via Convex Optimization (*AMICO*)^128^ framework to fit NODDI with fixed parallel intrinsic and isotropic diffusivities of 0.0011 and 0.003 mm²/s, respectively^129^.

#### Diffusion kurtosis imaging

For sensitivity analyses (see below), we also calculated tract-wise summary measures of diffusion kurtosis imaging (DKI) metrics from *DIPY*^130^ version 1.8.0. The diffusion tensor imaging (DTI) model characterizes the 3D Gaussian diffusion process in each voxel by estimating a second-order tensor from the diffusion-weighted signal. In contrast, the diffusion kurtosis imaging (DKI) model extends the tensor formulation by quantifying the degree to which water diffusion deviates from Gaussian behavior. This is achieved by additionally modeling a fourth-order kurtosis tensor that captures microstructural complexity not represented by DTI. Estimates of DTI parameters from a DKI model may be more reliable than from a standalone DTI model^131^. DKI scalar maps included axial diffusion (AD), fractional anisotropy (FA), mean diffusion (MD), radial diffusion (RD), kurtosis fractional anisotropy (KFA), mean kurtosis (MK), mean kurtosis tensor (MKT), axial kurtosis (AK), and radial kurtosis (RK).

### Tractography and tractometry

To delineate person-specific WM tracts, we used DSI Studio’s AutoTrack, which yielded 62 major WM tracts per participant (**Figure 1**). We calculated tract-specific microstructure measures (as described above) which we took the median of within each tract (**Figure 1**). In addition, we quantified 18 tract-level macrostructural properties^59^, including total volume, the volume of each length-wise quarter, tract length and span, surface area, trunk geometry (curl, elongation, and irregularity), as well as the surface area, volume, and radius of each end region (**Figure 1**). For the primary analysis, the feature set consisted of three NODDI-derived microstructural measures (median value within a tract) and 18 macrostructural measures per WM tract, yielding 21 metrics across 62 tracts (1,302 tract-wise imaging features per participant). In sensitivity analyses, we instead used nine DKI-derived microstructural measures together with 18 macrostructural measures across the same 62 tracts, resulting in 1,674 tract-wise imaging features per participant.

### Image Quality Measurement

To control for variation in scan quality, we used a previously-validated dMRI quality control (QC) measure T1 post DWI contrast as described in Meisler et al., 2026^58^.

### Quantification of the Exposome

To characterize environmental exposures, we used a previously validated exposome score (“general exposome”)^52^. In brief, this score was constructed using exploratory structural equation modeling applied to 354 environmental measures, followed by confirmatory bifactor modeling^52^. These 354 measures included individual-level exposures (e.g., youth-and caregiver-reported measures) and geocoded neighborhood-level indicators. This procedure yielded one general exposome factor and six orthogonal subfactors capturing distinct environmental domains: School, Family Values, Family Turmoil, Dense Urban Poverty, Extracurriculars, and Screen Time^52^. The general factor (general exposome) primarily reflects family-and neighborhood-level socioeconomic status (SES), with higher scores corresponding to more advantaged socioeconomic environments. **Supplementary Table 2** shows how the exposome scores were derived, including which environmental variables were included along with factor loadings, as described in the original manuscript.

### Quantification of Cognition

To characterize cognition, we used a previously validated general cognition score derived from principal component analysis of nine neurocognitive assessments collected at baseline in the ABCD study^132^. In this prior work, Bayesian probabilistic principal component analysis identified three broad cognitive domains: general cognitive ability, executive function, and learning/memory. Here, we focus specifically on the general cognitive ability component, which accounted for approximately 21% of the variance in the original neurocognitive battery and loaded most strongly on measures of crystallized verbal ability, including the NIH Toolbox Picture Vocabulary and Oral Reading tests^132^.

### Feature Harmonization

To reduce site-related variability in imaging features, we harmonized all tract-wise measures across sites using CovBat^68^, which extends ComBat^133^ capabilities by additionally preserving batch-wise covariance structures. Batch was set to a variable that accounts for the site, machine, and machine software associated with a scan. The CovBat model included protected covariates (age, sex, the general exposome score, and the orthogonal exposome subfactor scores), so that variance tied to these variables was retained after harmonization. In analyses that included cognition, we additionally protected baseline neurocognitive scores. To avoid information leakage between discovery and replication analyses, harmonization was performed using split-specific training. Specifically, we fit CovBat parameters using the discovery split as the training set and applied the learned harmonization to all participants (including the replication split). We then repeated the procedure with the replication split as the training set and applied it to all participants, generating a second harmonized dataset for the second half of the data. This procedure ensures that there is no information leakage between the discovery and replication samples.

### Mass-Univariate Models

We tested tract-wise associations with the exposome using ordinary least squares (OLS) regression. For each tract-wise imaging feature (1,302 total), we modeled the imaging feature as a function of the general exposome score, while adjusting for age, sex, and image quality^58^. We also included the orthogonal exposome subfactors as covariates (School, Family Values, Family Turmoil, Dense Urban Poverty, Extracurriculars, and Screen Time). Outcomes and all continuous predictors were *z*-scored prior to fitting.

We summarized association strength using the incremental variance explained by the general exposome term (ΔR^2^), defined as the change in model fit when adding the exposome to a reduced model excluding it. For visualization, we converted this quantity to a signed effect size by taking the square root of ΔR^2^ and assigning the sign of the exposome regression coefficient, yielding a signed semi-partial correlation (i.e., √ΔR²), which reflects the unique contribution of the exposome to each feature. The significance of the association between an imaging feature and the exposome was assessed using analysis of variance (ANOVA) comparing the full and reduced models. All *p*-values were corrected for multiple comparisons across tract-wise tests using the Benjamini-Hochberg false discovery rate (FDR) procedure^134^. Analyses were conducted separately in the two split half samples (discovery/replication) to assess robustness of effects across independent subsets. We collated these effect sizes from each sample into a vector and split half concordance was quantified by the Pearson correlation between effect size vectors.

### Alignment of White Matter-Exposome Associations with Cortical Hierarchies

Next, we sought to place our results in the context of the cortical hierarchy. To do this, we used principal component analysis (PCA) to summarize the dominant patterns in tract-wise exposome association profiles and tested whether tract scores aligned with the sensorimotor-association (S-A) axis^135^. Tract-wise association profiles were summarized using PCA applied to tract-by-metric matrices of signed √ΔR^2^ effect size values (semi-partial correlations). Separate matrices were constructed for the discovery and replication splits, with rows corresponding to tracts and columns corresponding to metrics. Effect sizes were then averaged across splits to form a single tract-by-metric matrix, which was z-scored across tracts to place metrics on a comparable scale. PCA was then fitted to the standardized data, which yielded tract scores and metric loadings.

To test whether the resulting components aligned with known cortical hierarchies, we related tract principal component (PC) scores to the cortical hierarchy defined by the S-A axis^135^. Using a validated tract-to-cortex mapping from DSI Studio^136^, we identified the subset of overlapping tracts between datasets (*n*=22, all association tracts), defined their connected cortical regions, and assigned each cortical region an S-A axis score^135^. Then, as in previous work^15^, we computed the S-A axis range for each tract as the difference between the maximum and minimum S-A scores across its connected cortical regions, reflecting the extent to which the tract spans the cortical hierarchy. We then tested the association between tract S-A axis range and the first PC score across tracts using Pearson’s correlation coefficient (*r*). Statistical significance was evaluated using permutation testing with 10,000 permutations of the S-A axis range value for each tract.

### Multivariate Prediction of Exposome Scores from White Matter Features in Unseen Data

We used split half multivariate ridge regression to evaluate the ability of the complex pattern of WM features to predict the exposome scores of an unseen individual. Analyses used the demographically matched split halves of the ABCD sample to evaluate out-of-sample performance. Models were trained in one split half and tested in the other, and then the direction was reversed.

We conducted analyses using three feature sets: microstructure-only, macrostructure-only, and a combined microstructure-plus-macrostructure feature set. Exposome scores were predicted from tract-wise imaging features using ridge regression, with the regularization parameter selected via nested cross-validation on the training data.

Prior to model fitting, nuisance covariates (age, sex, and image quality) were regressed from imaging features using a linear model fit on the training data only and applied to both training and test sets. Residualized imaging features and exposome scores were then standardized using scaling parameters estimated from the training data and applied to the held-out test data to avoid information leakage.

Model performance was quantified in the held-out split using Pearson’s correlation coefficient (*r*) between observed and predicted exposome scores. Statistical significance was assessed with permutation testing. Training labels were randomly permuted (*n*=1,000 permutations) and the full train-test pipeline was repeated to generate a null distribution of out-of-sample *r* values. Empirical *p*-values were computed as the proportion of null *r* values exceeding the observed *r* value.

To better interpret the multivariate models, we applied a Haufe transformation to the ridge regression feature weights. Haufe-transformed weights were calculated separately for each split half. We then compared these multivariate Haufe-transformed weights to the mass-univariate partial correlations using a Pearson correlation.

## Sensitivity Analyses

We conducted multiple sensitivity analyses to assess robustness. First, we repeated the primary analyses using diffusion kurtosis imaging (DKI) microstructure metrics in place of NODDI microstructure measures. Second, we evaluated the impact of global brain size by repeating models with estimated brain volume included as an additional covariate. Third, we tested an alternative multivariate modeling approach by fitting partial least squares (PLS) regression models in place of ridge regression. As with the main analysis using ridge regression, preprocessing, nuisance regression, scaling, and component selection were performed exclusively within the training data to prevent leakage into the test set. The number of PLS components was selected using five-fold nested cross-validation within the training data, maximizing the Pearson correlation between predicted and observed outcomes. Fourth, we compared the effect size of the multivariate association with the general exposome to alternative measures such as parental education and household income. We quantified the incremental predictive value of the exposome beyond these measures by calculating signed √ΔR² effect size values (semi-partial correlations; as described in the *Mass-Univariate Modeling* section). Fifth and finally, we tested sex-by-exposome interaction effects in mass-univariate models to assess whether exposome-WM associations differed by sex.

### Associations with Cognition Beyond the Exposome

To determine whether WM-cognition associations reflect variance shared with the exposome, we conducted both mass-univariate and multivariate predictive analyses. First, we performed mass-univariate analyses (see procedure in *Mass-Univariate Modeling* subsection) and quantified tract-wise associations with cognition. We then compared effect size estimates for cognition with and without adjustment for the exposome to assess how tract-level associations changed after accounting for shared variance with the exposome. We then tested whether tract-wise WM features predict cognition in held-out participants and compared the ability of WM features to predict cognition above and beyond the exposome. Using the same split half ridge regression framework (see procedure in *Multivariate Prediction* subsection), we predicted cognition from tract-wise imaging features and evaluated performance on held out data using Pearson’s correlation between observed and predicted cognition scores. To assess overlap with the exposome, we reran these cognition prediction models with and without including the exposome score as a nuisance covariate. Analogously, we evaluated the degree to which WM predicted the exposome score above and beyond cognition. To do this, we fit models predicting the exposome from tract-wise features with and without covarying for cognition.

### Evaluating Generalizability in HBN

#### HBN sample

To assess robustness and generalizability of WM-exposome associations, we conducted external replication analyses in the Healthy Brain Network (HBN)^63^. Importantly, HBN differs substantially from the ABCD study in sample ascertainment, clinical enrichment, construction of the exposome measure, and imaging acquisition parameters^63^. This makes HBN a rigorous test of the generalizability of our results. HBN is a self-referred study of children and adolescents recruited from the New York City area (ages 5–22)^63^. Parents or participants typically self-refer the study due to mental health or learning concerns, resulting in higher levels of psychopathology in the sample compared to the ABCD study^63^.

#### MRI Acquisition

This study analyzes 3T MRI data from the HBN, which were acquired at three sites (MRI derivatives were harmonized across sites as described below). Data were collected at the Rutgers University Brain Imaging Center on a 3T Siemens Tim Trio scanner as well as at the CitiGroup Cornell Brain Imaging Center and the CUNY Advanced Science Research Center on 3T Siemens Prisma Scanners. T1w images were acquired with an MPRAGE sequence with the following parameters: repetition time of 2,500 ms, echo time of 3.15 ms, inversion time of 1,060 ms, flip angle of 8 degrees, 224 slices and a voxel resolution of 0.8 mm isotropic. Diffusion scans were acquired with a multiband factor of three in two shells of bLJ=LJ1,000 and 2,000 s/mm2 in the A–P phase-encoding direction, with 72 slices and multiband acceleration = 3. In total, 64 directions were acquired per shell (128 directions total) along with 1 b = 0 volume. The following parameters were used for the diffusion acquisition: repetition time of 3,320 ms, echo time of 100.2 ms and a voxel resolution of 1.8 mm isotropic. A reverse phase-encoding b = 0 was additionally acquired for use as an echo planar image (EPI)-based field map in susceptibility distortion correction.

#### Diffusion MRI Preprocessing

Diffusion MRI preprocessing and reconstruction for HBN followed the same general QSIPrep and QSIRecon-based pipeline described for ABCD. Here, we summarize dataset-specific details that differ. Diffusion MRI data were preprocessed using QSIPrep 1.0.2^66^, based on Nipype 1.9.1^107,108^. Data were postprocessed using QSIRecon 1.1.1^137^. Preprocessed diffusion data were resampled to 2 mm isotropic resolution (rather than preserving the native 1.7 mm resolution). Preprocessing configuration did not incorporate TOPUP-based susceptibility distortion correction. Eddy current and motion correction included slice-wise outlier detection and imputation procedures specific to this dataset. A quadratic first level model was used to characterize Eddy current-related spatial distortion. q-space coordinates were forcefully assigned to shells. Field offset was attempted to be separated from subject movement. Shells were aligned post-eddy. Eddy’s outlier replacement was run^125^. Data were grouped by slice, only including values from slices determined to contain at least 250 intracerebral voxels. Groups deviating by more than 4 standard deviations from the prediction had their data replaced with imputed values. Final interpolation was performed using the jac method. For NODDI reconstruction, the AMICO implementation was used with a parallel intrinsic diffusivity of 0.0017 mm²/s (isotropic diffusivity = 0.003 mm²/s), differing from the value used in ABCD. All other preprocessing, reconstruction, and tractography steps were consistent with the ABCD workflow.

#### General Exposome Calculation

As previously described, the general exposome score in HBN reflects neighborhood-level socioeconomic context and was derived from census and environmental protection agency data using exploratory structural equation modeling with a bifactor structure^86,92^. Participants were assigned scores based on their residential census block group^86,92^. Thus, unlike ABCD, where the exposome incorporates both individual-level (self-and parent-reported) and geocoded environmental measures, the HBN exposome is derived exclusively from 43 geocoded neighborhood-level indicators (see **Supplementary Table 3** for top loadings).

#### Sample Construction

External replication analyses were conducted in the subset of HBN participants with available diffusion MRI and exposome data (*n*=869). For this subset of participants, we derived tract-wise microstructural NODDI metrics (ICVF, OD, ISOVF) and tract-level macrostructural measures using the same tract definitions as in ABCD. For HBN, as in prior work^86,138^, we define image quality as the QSIPrep-derived T1 neighbor correlation metric. Analyses were restricted to tract-wise features present in both ABCD and HBN.

#### Harmonization to ABCD and nuisance regression

To reduce site effects across studies, we harmonized HBN features to the ABCD data using CovBat-GAM, using the largest ABCD site with a Siemens scanner as the reference (i.e., site 16). We selected CovBat-GAM because HBN spans a wider age range (5-22 years), allowing us to account for nonlinear age effects across this range. The model protected variance associated with age, sex, and the exposome. We saved this harmonized HBN dataset for univariate analyses.

#### Mass-univariate replication of exposome-white matter associations in HBN

We then residualized tract-wise features for age (using a natural cubic spline, *df*=3), sex, and t1 neighborhood correlation, producing a harmonized and residualized HBN feature set for the multivariate analyses. We tested for tract-wise associations with the general exposome within HBN using the same mass-univariate framework as in ABCD (see procedure in *Mass-Univariate Modeling* subsection). For each tract-wise feature, we fit a Generalized Additive Model (GAM) with general exposome as the predictor and adjusted for age, sex, and image quality, with a smooth term for age. Association strength was summarized using signed √ΔR² effect size values (semi-partial correlations) for the general exposome term. We then correlated these tract-wise effect size estimates to the corresponding estimates obtained in ABCD, yielding a measure of concordance in association strength across studies.

#### Multivariate Prediction of Exposome Scores from White Matter Features in Unique Dataset

We evaluated out-of-sample generalization from ABCD to HBN by training multivariate ridge regression models in the split half ABCD samples (see procedure in *Multivariate Prediction* subsection) and applying them to unseen HBN participants without further tuning. Models trained separately in each of the matched halves of the ABCD data were used to predict unseen general exposome scores in HBN, yielding two independent external performance estimates. Performance was quantified using Pearson correlation (*r*) between observed and predicted general exposome scores in HBN. Significance was assessed with permutation testing in which ABCD training labels were permuted to generate a null distribution of *r* values (*n*=1,000 permutations).

## Resource availability

All code used for data analysis and figure generation is publicly available on the project’s GitHub repository: https://github.com/PennLINC/macedo_wm_exposome. A detailed replication guide is available here: https://pennlinc.github.io/macedo_wm_exposome/.

Data used in the preparation of this article were obtained from the *Adolescent Brain Cognitive Development*JJ (*ABCD*) Study, held in the NIH Brain Development Cohorts Data Sharing Platform. This is a multisite, longitudinal study designed to recruit more than 10,000 children age 9–10 and follow them over 10 years into early adulthood. The ABCD dataset grows and changes over time. The raw ABCD data used in this report came from ABCD release 6.0: https://doi.org/10.82525/jy7n-g441. Data from the Healthy Brain Network is accessible through https://fcon1000.projects.nitrc.org/indi/cmihealthybrainnetwork/.

## Acknowledgements

This research was supported by funding from the National Institutes of Health (R37MH125829 to D.A.F. and T.D.S.; T32MH019112 to S.L.M.; 2R01MH112847 to R.T.S. and T.D.S.; R01MH120482 to T.D.S.; 2R01MH113550 to T.D.S.; R01MH123550 to R.T.S.; F30MH138048 to K.Y.S.; RF1MH121868, RF1MH121867, RF1MH126699, R01AG060942, U19AG066567, R01EY033628, and R01EB027585 to A.R.; R01MH134886 to R.B.; T32MH016804 and T32MH018951 to V.J.S.; R01MH133843 to A.A.B.; F31MH136685 to J.B. Additionally, S.L.M. was supported by the Hartwell Foundation. G.S. was supported by a postdoctoral fellowship from the Canadian Institutes of Health Research (CIHR). A.S.K. is supported by a NARSAD Young Investigator Award from the Brain and Behavior Research Foundation. M.D.H. was supported by the German Research Foundation (project number 572317568). LMS was supported by an NSF SBE Postdoctoral Research Fellowship (#2507497).

The ABCD Study® is supported by the National Institutes of Health and additional federal partners under award numbers U01DA041048, U01DA050989, U01DA051016, U01DA041022, U01DA051018, U01DA051037, U01DA050987, U01DA041174, U01DA041106, U01DA041117, U01DA041028, U01DA041134, U01DA050988, U01DA051039, U01DA041156, U01DA041025, U01DA041120, U01DA051038, U01DA041148, U01DA041093, U01DA041089, U24DA041123, U24DA041147. A full list of supporters is available at https://abcdstudy.org/about/federal-partners/.

Thank you also to Deanna M. Barch, who provided valuable feedback on this manuscript.

## Author contributions

Conceptualization: B.M.; Software: B.M., S.L.M., M.C., and F.-C.Y.; Data curation: S.L.M., M.C., D.A.F., R.B., T.M.; Formal analysis: B.M.; Visualization: B.M., with feedback from J.B., S.L.M., L.S., V.J.S., A.K., and T.D.S.; Resources: V.J.S., T.M., A.S.K.; Validation: J.B.; Funding acquisition: T.D.S.; Methodology: B.M., T.D.S., J.B., S.L.M., L.S., M.H., A.A.B., R.B., T.M.; Supervision: T.D.S. and A.A.B.; Writing – original draft: B.M.; Writing – review & editing: J.B., S.L.M., L.S., M.H., A.P.M. R.B., T.M., and T.D.S.; All authors approved the manuscript.

## Declaration of generative AI and AI-assisted technologies

All draft text was initially written by the authors. During the revision of this work, the authors used ChatGPT to offer suggestions on how to improve the readability and flow of the text. After using this tool/service, the authors further edited and rewrote the text and take full responsibility for all of the content of the article.

