## supplemental materials for "White matter reflects the childhood exposome"

**Supplementary Materials**

This Supplementary Material provides additional analyses, robustness checks, and methodological details supporting the main findings.

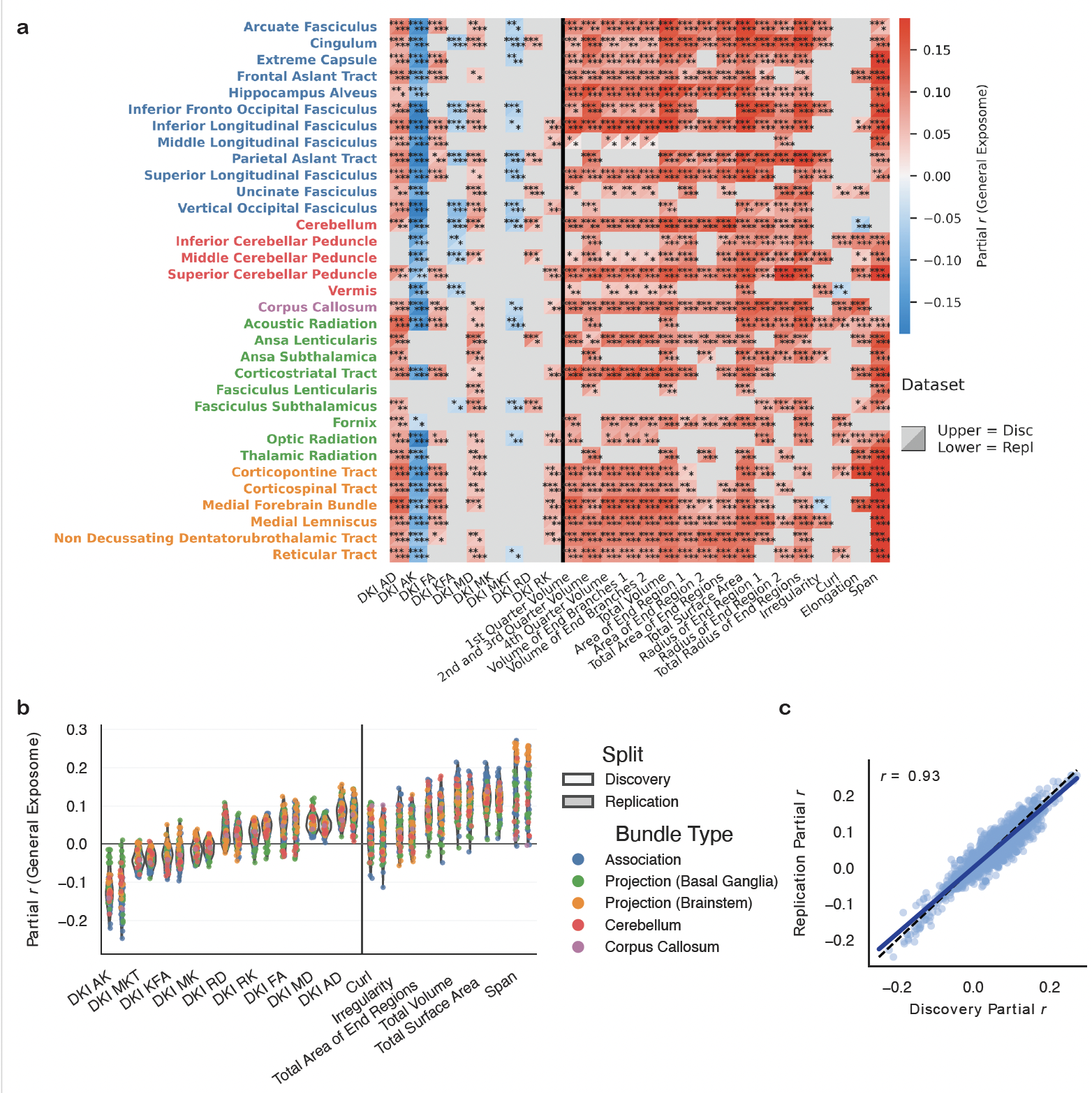

**Supplementary Figure 1: Mass-univariate associations between white matter micro- and macrostructure and the exposome using DKI metrics.** Same analyses and visualization as in **Figure 2** but using nine diffusion kurtosis imaging (DKI) microstructural metrics instead of NODDI metrics. Panels are as described in **Figure 2**.

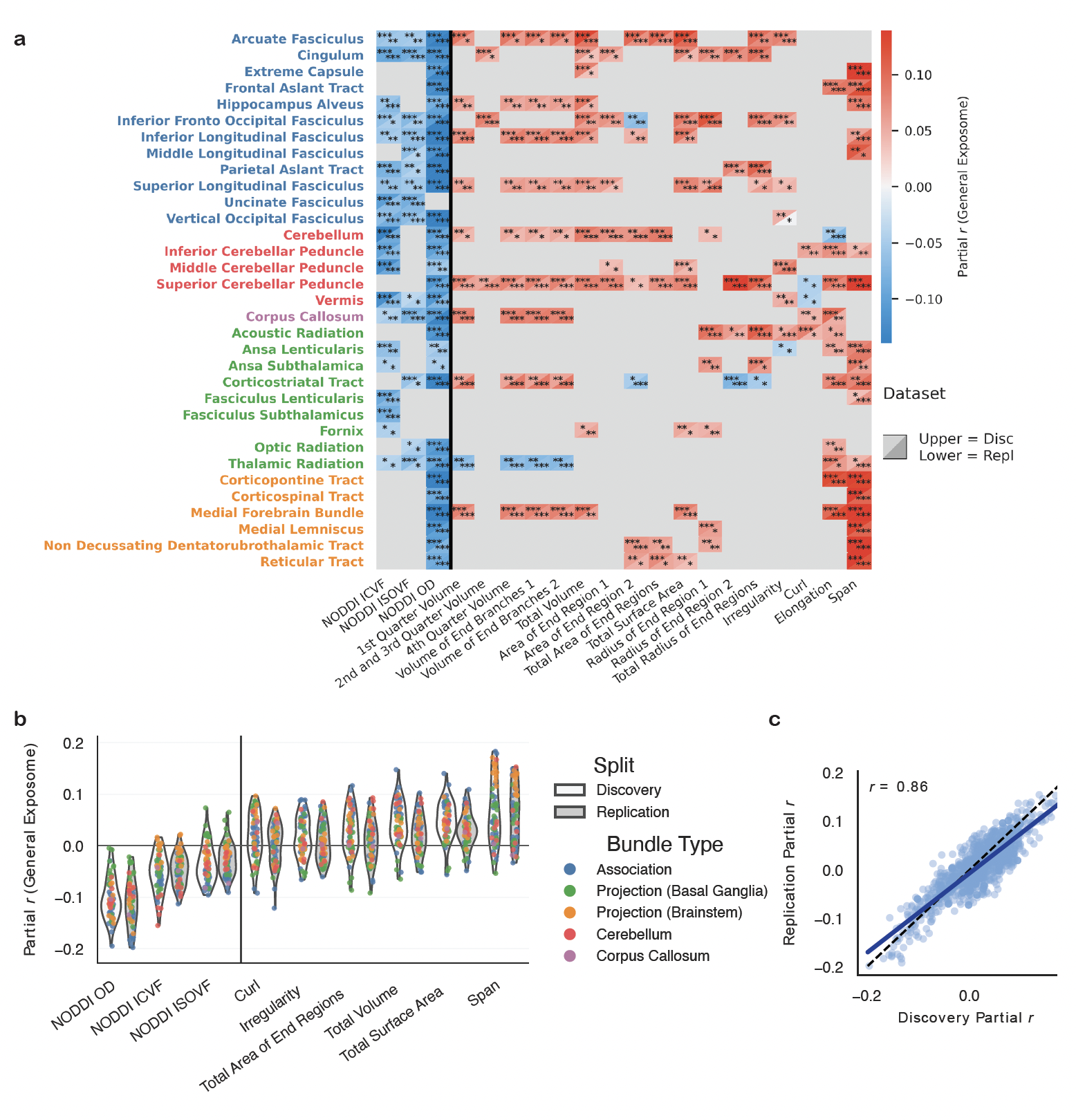

**Supplementary Figure 2: Mass-univariate associations between white matter micro- and macrostructure and the exposome with total brain size correction.** Same analyses and visualization as in Figure 2, but with all models additionally adjusted for total brain size prior to analysis. Panels are as described in Figure 2.

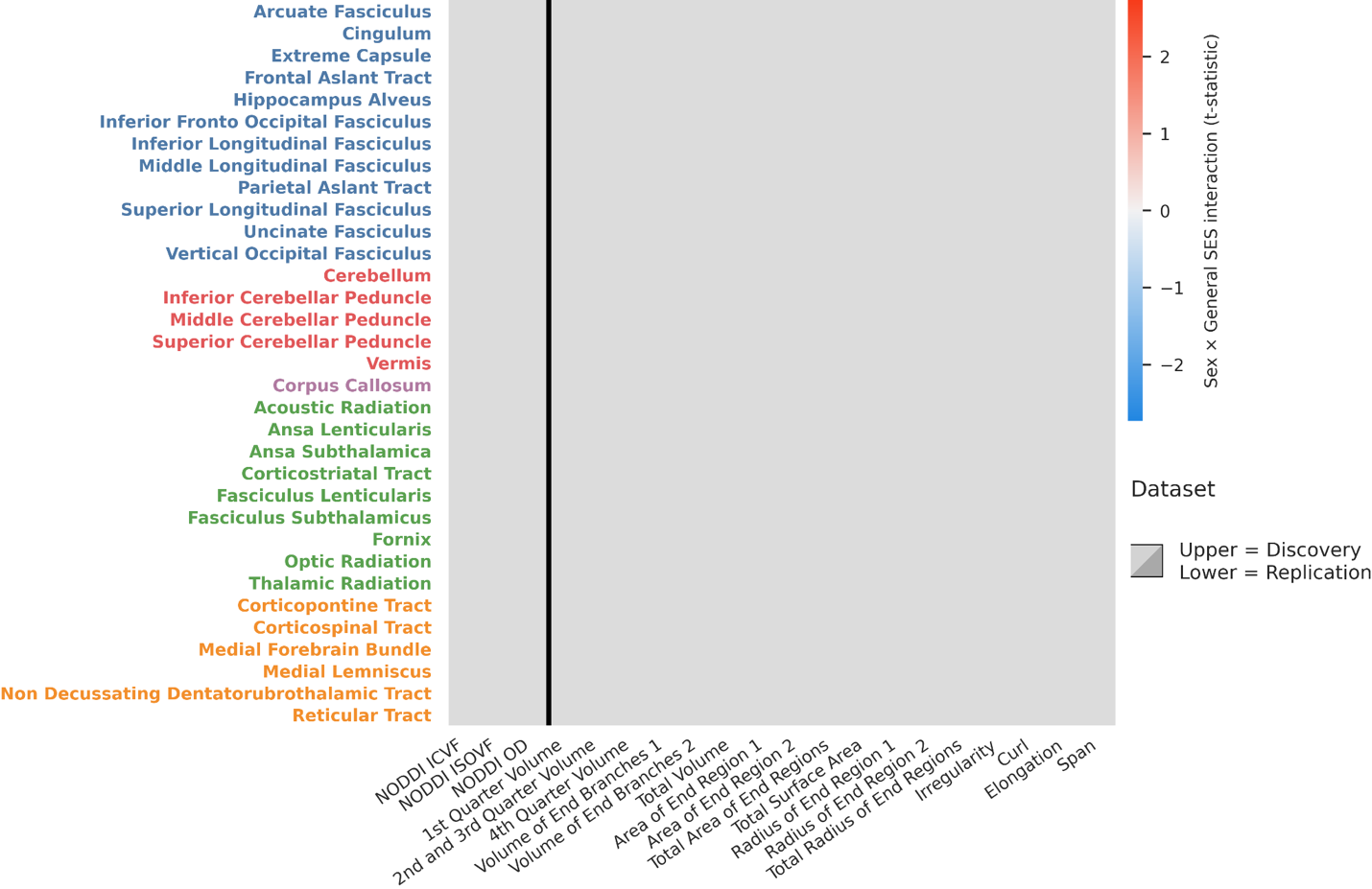

**Supplementary Figure 3: Sex-by-exposome interaction effects on white matter micro- and macrostructure.** Mass-univariate linear regressions including a sex × exposome interaction term, otherwise as in **Figure 2**. Heatmap displays t-statistics for the interaction term across all tract-wise microstructural and macrostructural metrics. No associations survived FDR correction.

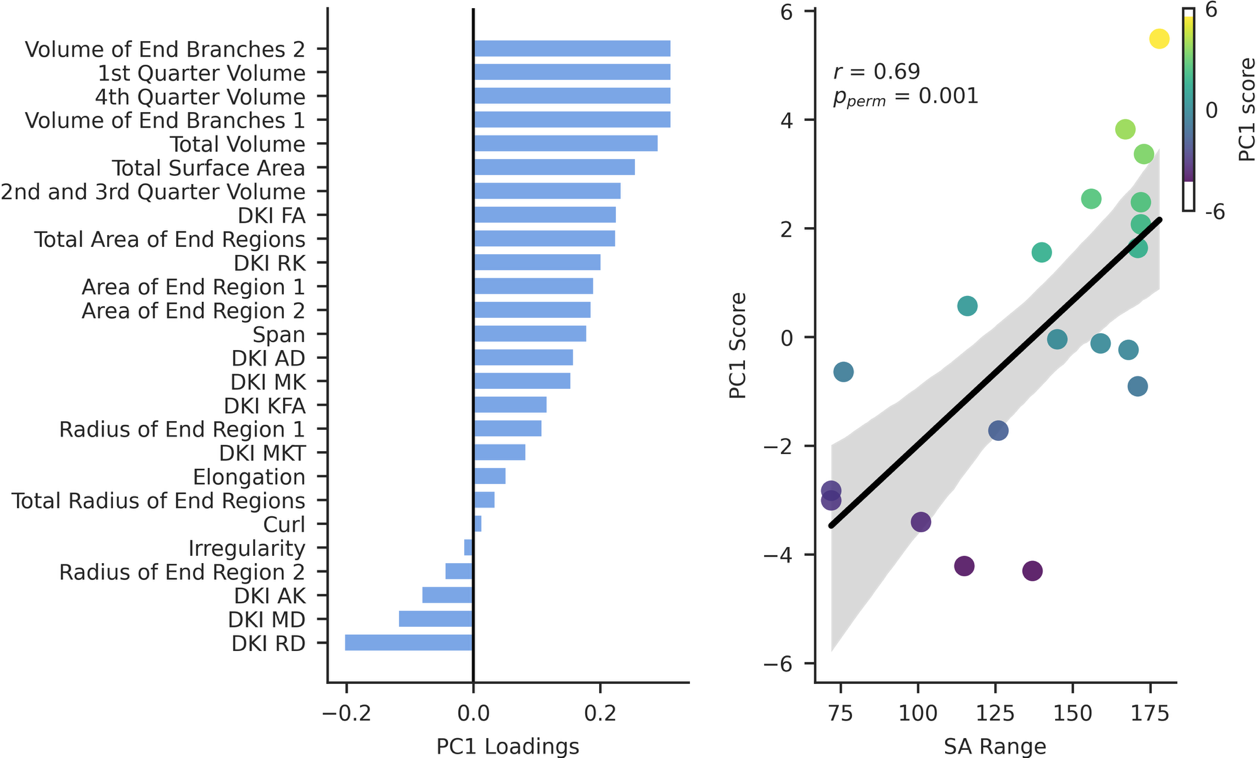

**Supplementary Figure 4: Principal components of white matter-exposome associations using DKI metrics.** Same analysis and visualization as in **Figure 3** but using DKI microstructural metrics instead of NODDI metrics. Panels are as described in **Figure 3**.

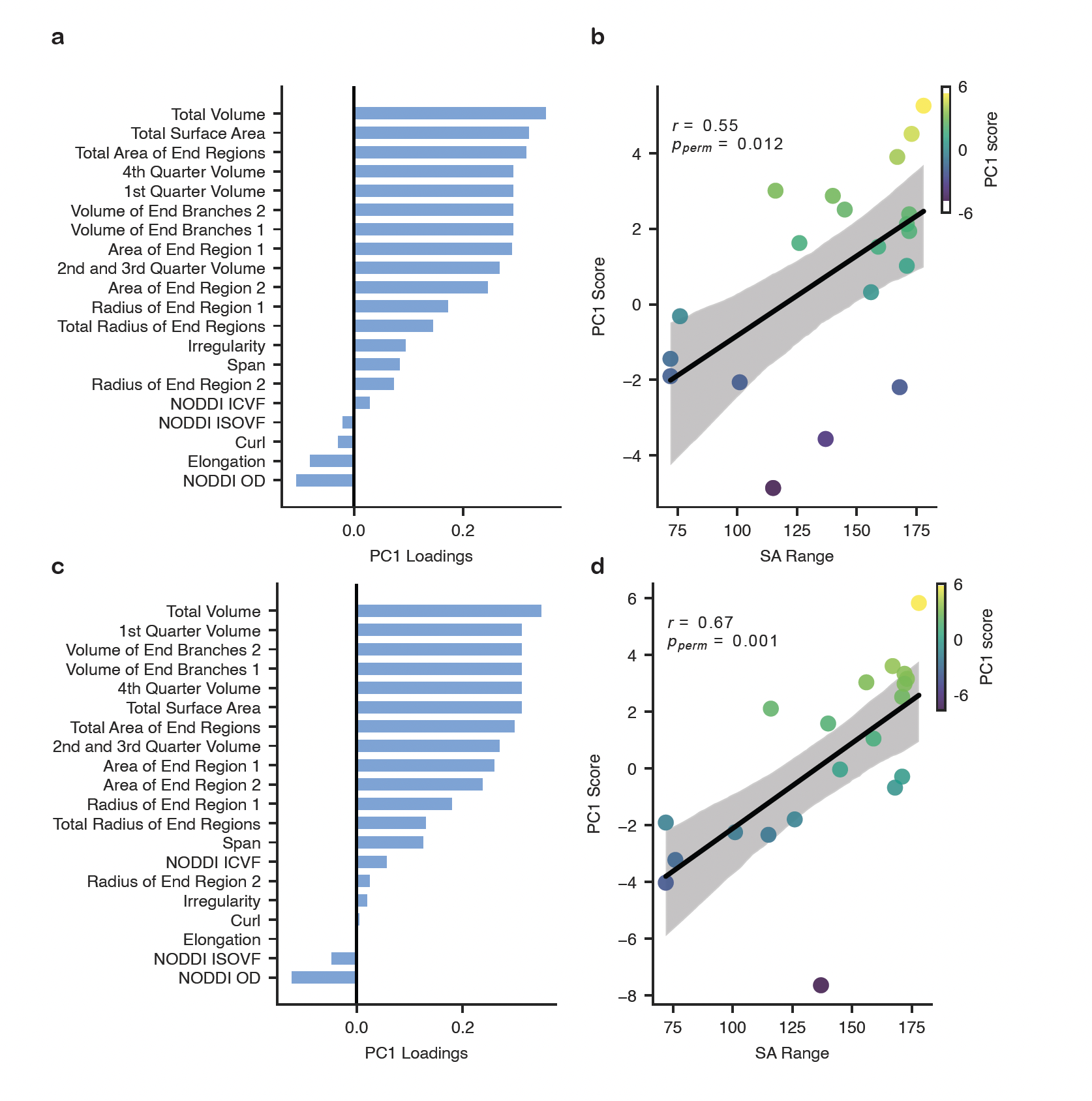

**Supplementary Figure 5: Principal components of white matter-exposome associations in split-half analyses.** Same analysis and visualization as in **Figure 3**, but performed separately in each split half rather than on averaged data. Panels are organized as in **Figure 3**: **a-b** show the discovery split and **c-d** show the replication split.

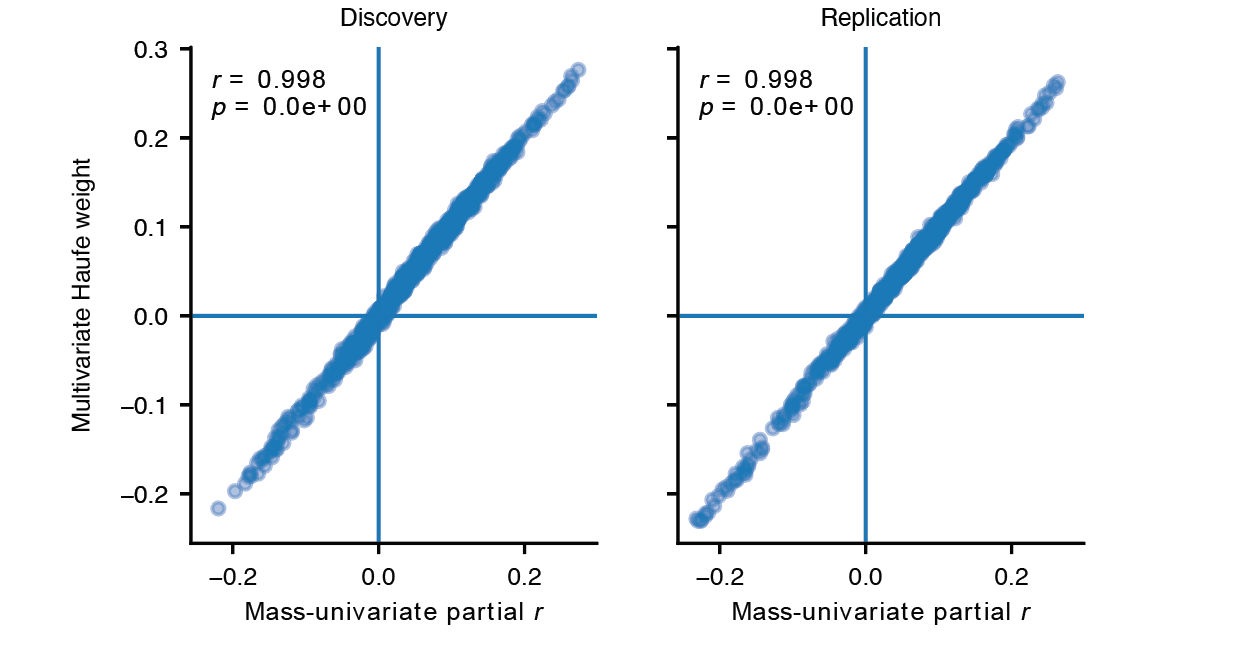

**Supplementary Figure 6**: **Multivariate Haufe-transformed weights align with univariate partial correlations.** Comparison of feature importance derived from multivariate ridge regression models and mass-univariate linear regression models. Haufe-transformed weights from the multivariate prediction models were correlated with tract-wise partial correlation (partial *r*) values from the mass-univariate models across imaging features. Each point represents a single tract-wise feature. Pearson’s correlation coefficients (*r*) quantify the correspondence between multivariate and univariate effect estimates, computed separately in each split-half sample.

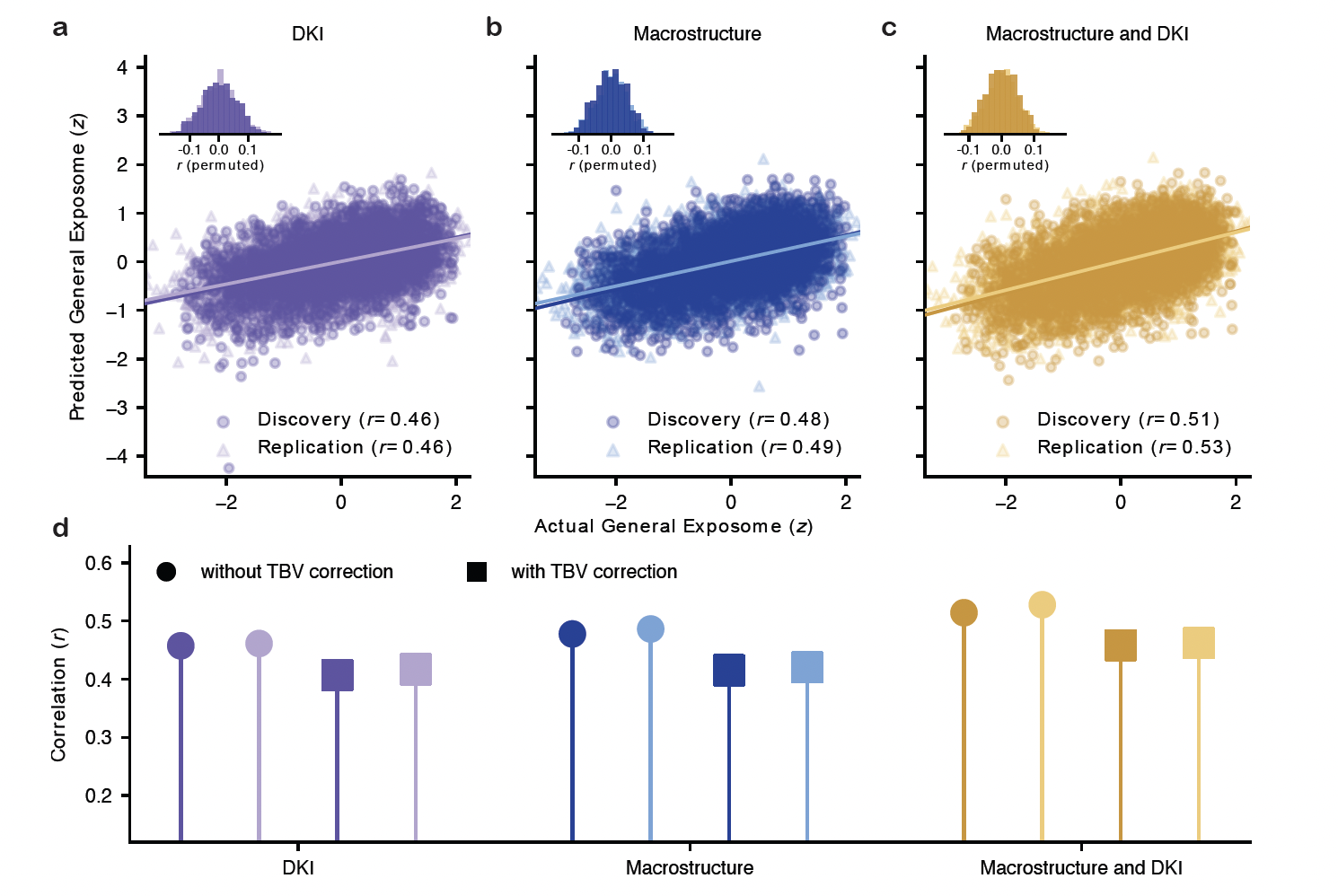

**Supplementary Figure 7: White matter features robustly predict childhood exposome in unseen data using DKI microstructural metrics.** Same analysis and visualization as in **Figure 4** but using DKI microstructural metrics instead of NODDI metrics. Panels are as described in **Figure 4**.

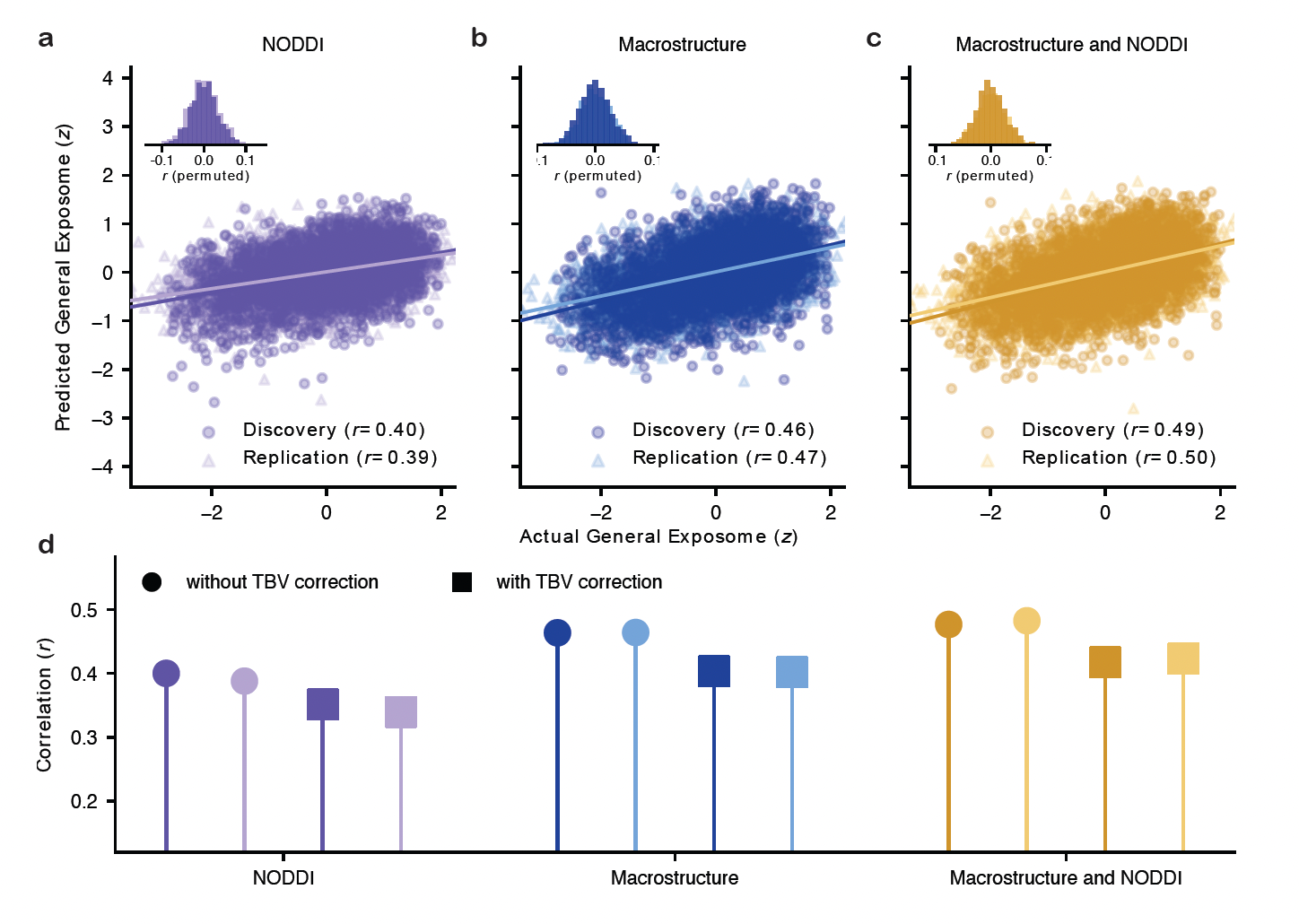

**Supplementary Figure 8: White matter features robustly predict childhood exposome in unseen data using partial least squares regression.** Same analysis and visualization as in **Figure 4** but using partial least squares regression instead of ridge regression. Panels are as described in **Figure 4**.

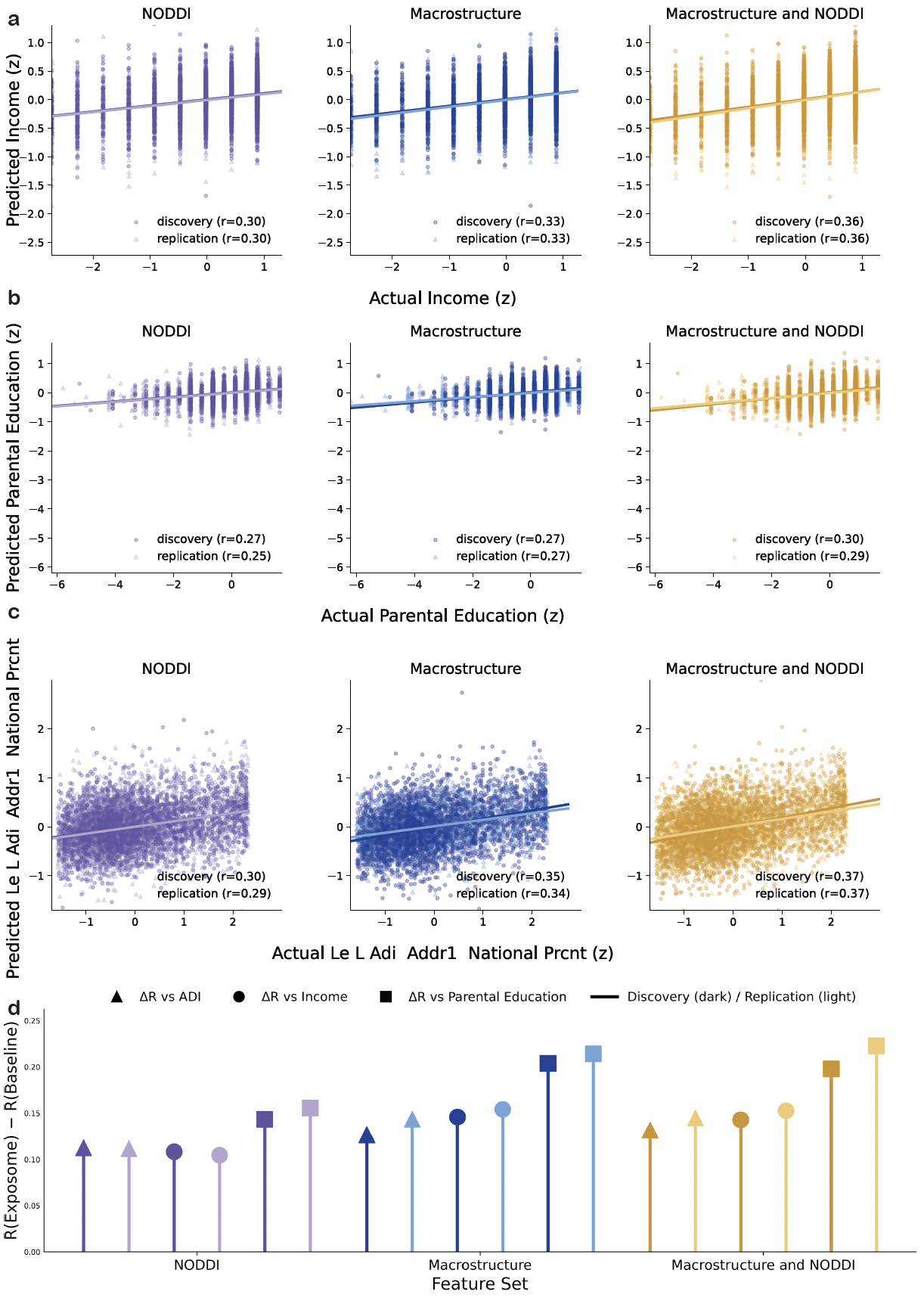

**Supplementary Figure 9: Prediction of socioeconomic indicators from white matter features.** Same analysis as in **Figure 4**, but with household income (**a**), parental education (**b**), area deprivation index (**c**), and used as outcome variables instead of the exposome score. Points show differences in predictive performance (ΔR), computed as the difference in out-of-sample Pearson correlation (*r*) between models predicting the exposome score and models predicting each socioeconomic indicator. Performance across different feature sets and split halves are shown in **d.**

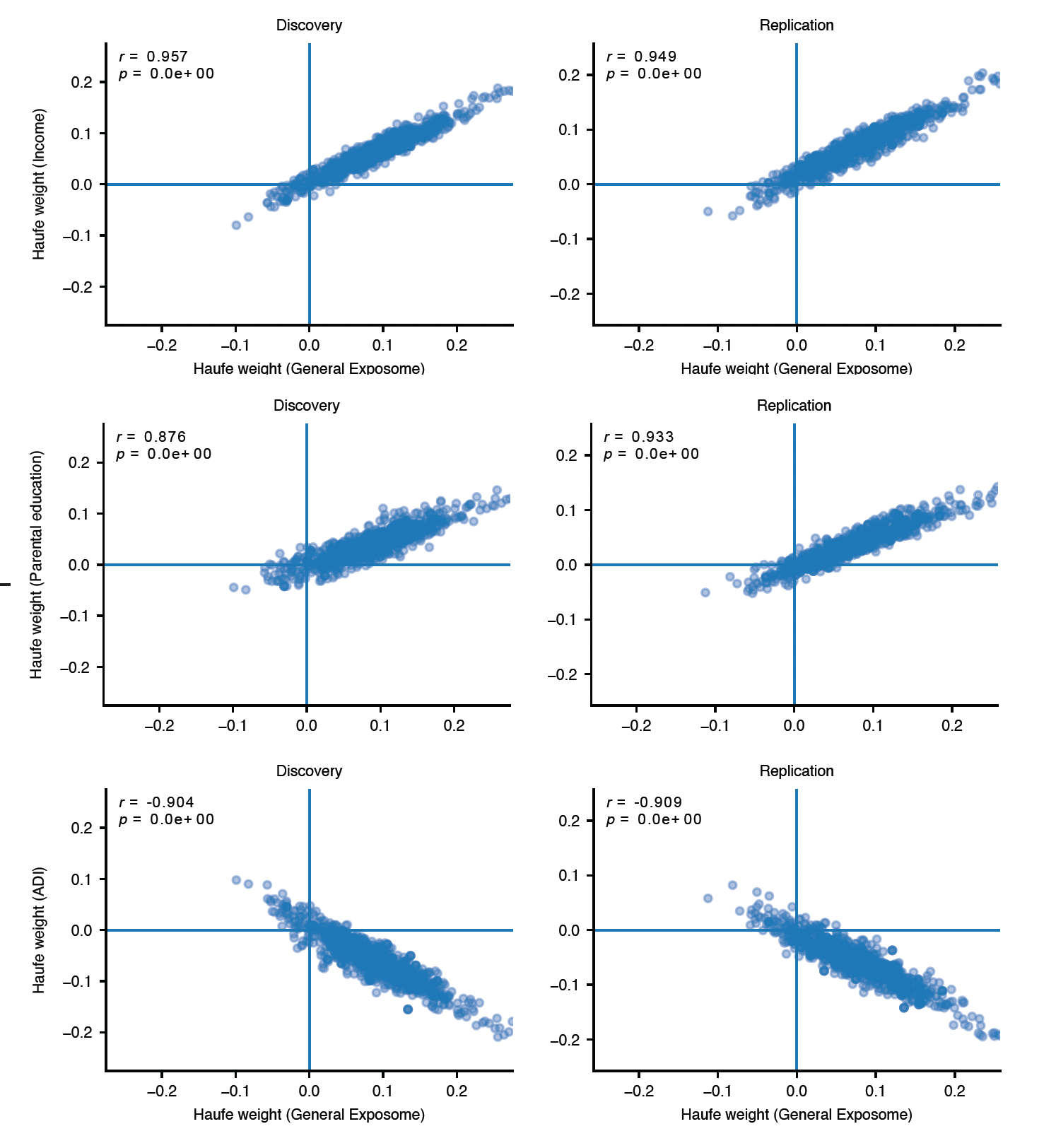

**Supplementary Figure 10: Concordance of multivariate feature weights across environmental outcome models.** Correlation between Haufe-transformed feature weights from models predicting the general exposome score and models predicting household income, parental education, and area deprivation index. Each point represents a tract-wise imaging feature. Results are shown separately for discovery and replication split halves. Pearson’s correlation coefficients (*r*) quantify the similarity of feature weights across outcome variables, indicating consistency of multivariate feature importance across related environmental indicators.

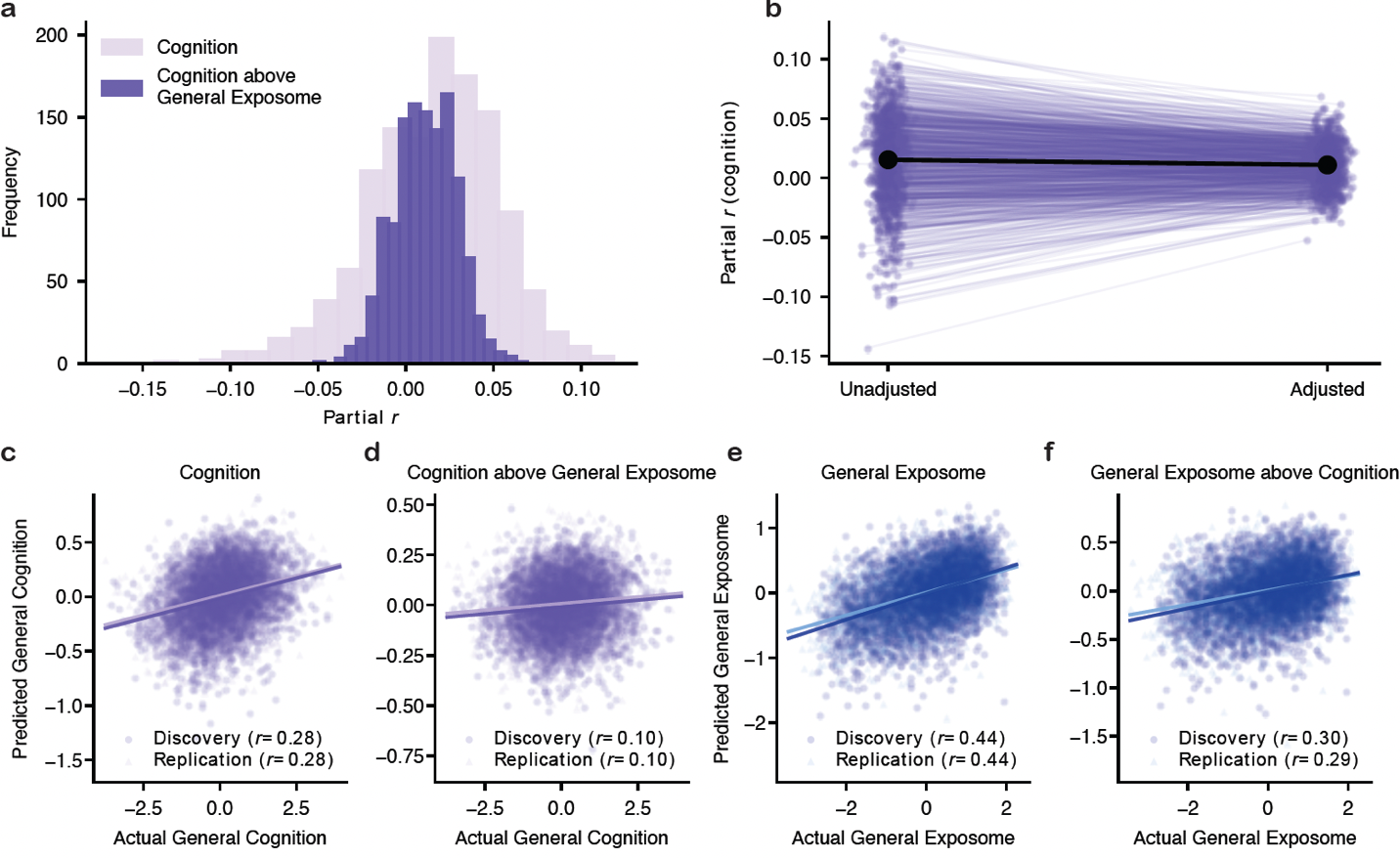

**Supplementary Figure 11: Adjustment for the exposome reduces cognition associations and prediction from white matter features after controlling for total brain size.** Same analysis and visualization as in **Figure 5**, but with all models additionally adjusted for total brain volume (TBV).

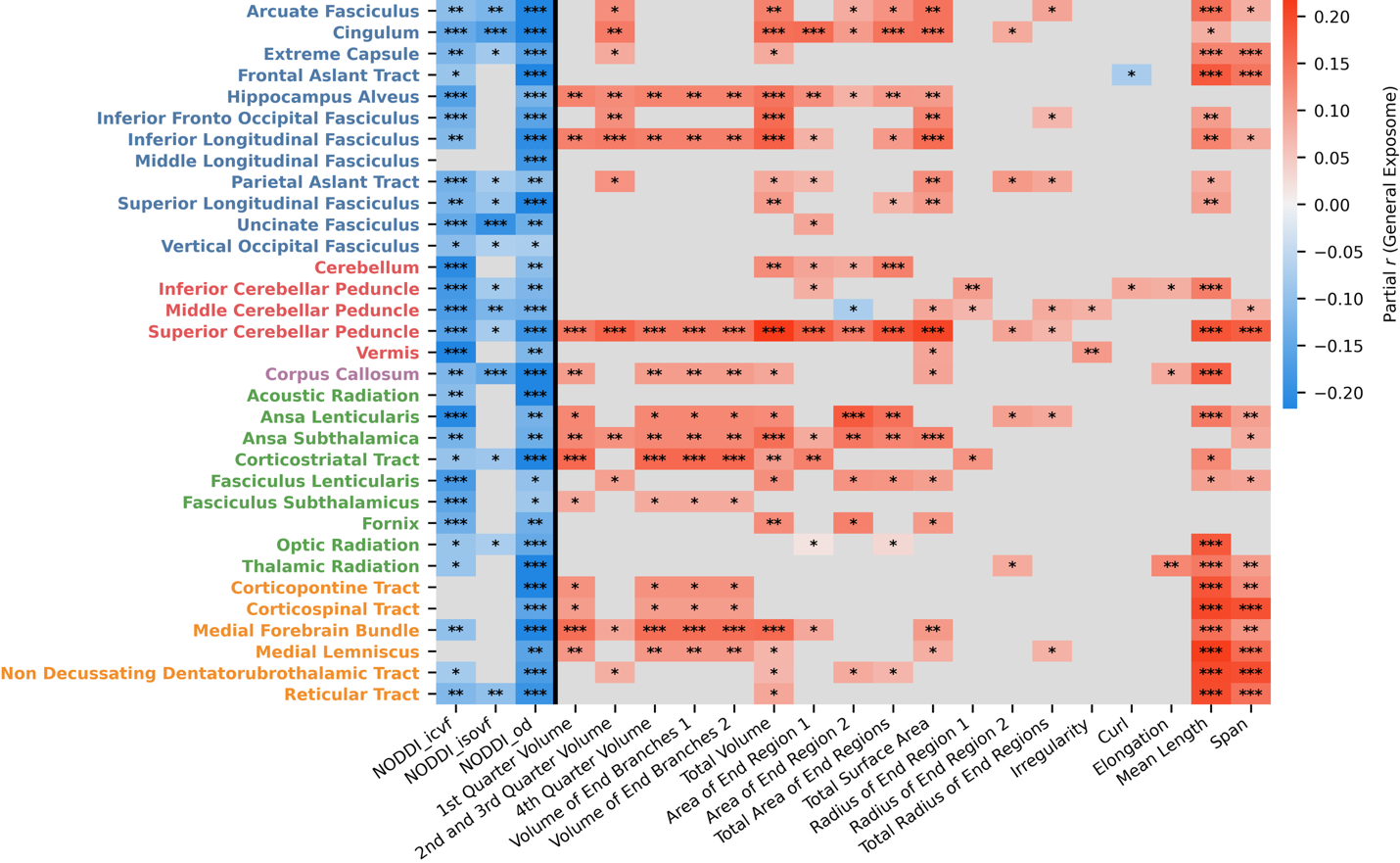

**Supplementary Figure 12: Mass-univariate associations between white matter features and the exposome in the Healthy Brain Network cohort.** Same analysis and visualization as in **Figure 2a** but conducted in the Healthy Brain Network (HBN) sample. Heatmap displays tract-wise effect sizes between WM microstructural and macrostructural features and the exposome score. Results are shown for HBN and correspond to the feature-wise associations compared with ABCD in **Figure 6a**.

| **Category** | **Metric** | **Definition** | **Interpretation** |
| --- | --- | --- | --- |
| Micro | Intracellular volume fraction (ICVF) | Fraction of tissue volume occupied by the intracellular neurite compartment | Higher values suggest greater neurite density |
| Micro | Isotropic volume fraction (ISOVF) | Fraction of isotropically diffusing water | Higher values suggest more free water |
| Micro | Orientation dispersion index (OD) | Degree of dispersion of neurite orientations | Higher values indicate greater fiber dispersion/fanning |
| Macro | Length (mm) | Average trajectory length of streamlines within a bundle | Overall tract extent |
| Macro | Span (mm) | Average endpoint-to-endpoint distance of bundle streamlines | Spatial reach of the tract |
| Macro | Diameter (mm) | Estimated bundle thickness derived from tract volume and length | Cross-sectional bundle size |
| Macro | Radius (mm) | Characteristic radius of the bundle end surface | Bundle caliber at the endpoints |
| Macro | Surface area (mm²) | Area of the voxelized tract surface | Extent of the tract boundary |
| Macro | Volume (mm³) | Total voxelized volume occupied by the bundle | Overall tract size |
| Macro | Trunk volume (mm³) | Voxelized volume of the tract core/trunk | Size of the main body of the tract |
| Macro | Curl | Ratio of length to span | Higher values indicate greater curvature/tortuosity |
| Macro | Elongation | Ratio of length to diameter | Higher values indicate a longer, thinner tract |

**Supplementary Table 1. Tract-level microstructural and macrostructural descriptors.** This table summarizes the NODDI-derived microstructural measures and tract geometry-based macrostructural measures used in the primary analyses. Microstructural measures were computed from AMICO-NODDI and summarized within each tract as the median value. Macrostructural measures were computed from tract geometry in DSI Studio and are based on the shape analysis framework described by Yeh [59].

| **Variable** | **Exposome Factor Loading** |
| --- | --- |
| Household Income | 0.78 |
| ABGD: Poverty | -0.695 |
| Parental Education | 0.68 |
| Parents Married | 0.572 |
| SAIQ: Miscellaneous Sports/Activities | 0.507 |
| SAIQ: Arts & Individual Sports | 0.475 |
| MACV: Religiosity | -0.465 |
| MACV: Family Image | -0.437 |
| ABGD: Traditional South/Midwest | -0.415 |
| Neighborhood Safety | 0.398 |
| ABGD: Crowding and Crime | -0.311 |
| FES: Family Conflict (Youth) | -0.271 |
| MACV: Independence | -0.247 |
| Screen Time | -0.235 |
| Substance Use Attitudes | 0.225 |
| SAIQ: Competitive Athletics | 0.223 |
| Parental Monitoring | 0.208 |
| Parental Rules on Substance Use | 0.201 |
| FES: Family Non-Conflict (Youth) | -0.182 |
| FES: Family Conflict (Parent) | -0.18 |
| MACV: Caring/Security | -0.147 |
| ABGD: Ozone | 0.122 |
| ABGD: Dense/Suburban | -0.121 |
| SRPF: Positive Feedback in School | -0.104 |
| SRPF: Enjoy School | 0.09 |
| Peer Deviance | -0.086 |
| TBIs | 0.085 |
| SRPF: Feel Involved in School | 0.075 |
| SRPF: Good Grades | -0.063 |
| ABGD: Air Pollution | -0.043 |
| FES: Family Non-Conflict (Parent) | 0.012 |
| ABGD: Retirement/Group Living | -0.009 |

**Supplementary Table 2:** Factor loadings from the longitudinal exploratory bifactor analysis for the general exposome factor in ABCD. Higher scores reflect more advantaged socioeconomic environments. Variables are ordered by absolute loading on the general exposome factor. Loadings are adapted from Keller 2024^52^. Full bifactor loadings, including subfactor-specific loadings, are reported in the original publication. Abbreviations: SRPF: School Risk and Protective Factors Survey; FES: Family Environment Scale - Family Conflict Subscale; MACV: Mexican American Cultural Values Scale; ABGD: Address-Based Geographic Data; SAIQ: Sports and Activities Involvement Questionnaire. All questionnaires were given to the full set of participants.

| **Variable** | **Exposome Factor Loading** |
| --- | --- |
| population density | 0.108 |
| public transit use | 0.144 |
| % married | **0.497** |
| vehicle traffic | 0.025 |
| hazardous waste proximity | 0.115 |
| respiratory hazard | 0.075 |
| median rooms per dwelling | **0.516** |
| % homeless | 0.045 |
| % veterans | -0.034 |
| median home value | **0.800** |
| % who walk to work | -0.031 |
| median owner upkeep cost | **0.799** |
| % in poverty | **-0.608** |
| housing newness | 0.152 |
| vehicles available | 0.275 |
| work-from-home ratio† | 0.358 |
| median rent | **0.776** |
| % abroad 1 year ago | 0.047 |
| professional:trade ratio‡ | 0.246 |
| soft:hard career ratio§ | -0.074 |
| % unemployed | -0.351 |
| owner:renter ratio | 0.236 |
| average household size | 0.096 |
| % with high school edu | **0.573** |
| % non-family households | -0.083 |
| % in graduate school | 0.149 |
| % without health insurance | **-0.489** |
| % same-sex couples | -0.162 |
| median family income | **0.887** |
| median age | 0.267 |
| % with children | 0.018 |
| % with retirement inc. | 0.121 |
| % widowed | -0.169 |
| % native English speakers | 0.141 |
| % with meals in rent | -0.010 |
| risk management facilities | -0.159 |
| airborne particulates | 0.006 |
| % mobile homes | -0.278 |
| superfund proximity | 0.156 |
| % vacant units | -0.279 |
| wastewater discharge | -0.018 |
| relative ozone conc. | 0.002 |
| % female | -0.031 |

**Supplementary Table 3:** HBN exposome loadings. Adapted from Moore 2016^92^ and Sydnor 2025^86^. Loadings with absolute value<0.30 removed for clarity unless it is the primary loading on a given factor; primary loadings on the general factor bolded; SES = socioeconomic status; inc = income; conc = concentration; †work-from-home ratio = total residents who work from home divided by total residence who travel to work in a car or truck; ‡professional:trade ratio = total residents who work in a "professional" field divided by total residents who work in a "trade" (see Moore 2016 for details); §soft:hard career ratio = total residents in a "soft" career divided by total residents in a "hard" career (see Moore 2016 for details); factor extraction = maximum likelihood; rotation = Jennrich-Bentler orthogonal bifactor.
